# Mosaic receptor-binding domain nanoparticles induce protective immunity against SARS-CoV-2 challenges

**DOI:** 10.1101/2022.02.18.480994

**Authors:** Dan Bi Lee, Hyojin Kim, Ju Hwan Jeong, Ui soon Jang, You Yeon Chang, Seokbeom Roh, Hyunbum Jeon, Eun Jeong Kim, Su Yeon Han, Jin Young Maeng, Stefan Magez, Magdalena Radwanska, Ji Young Mun, Hyun Sik Jun, Gyudo Lee, Min-Suk Song, Hye-Ra Lee, Mi Sook Chung, Yun Hee Baek, Kyung Hyun Kim

## Abstract

Recurrent spillovers of α- and β-coronaviruses (CoV) such as acute respiratory syndrome (SARS)-CoV, Middle East respiratory syndrome (MERS)-CoV, SARS-CoV-2, and possibly human CoV (NL63, 229E, OC43, and HKU1) have caused serious morbidity and mortality worldwide. Six receptor binding domains (RBDs) derived from α- and β-CoV that are considered to have originated from animals and cross-infected humans were linked to proliferating cell nuclear antigen (PCNA) heterotrimeric subunits, PCNA1, PCNA2, and PCNA3. These were used to form a scaffold-based mosaic multivalent antigen, 6RBD-np. Electron microscopic and atomic force microscopic images show a ring-shaped disk with six protruding RBDs, like jewels in a crown, with a size of 40 nm. Prime-boost immunizations with 6RBD-np in BALB/c mice elicited strong, dose-dependent antibody responses. In human angiotensin converting enzyme 2-transgenic mice, the same immunization induced full-protection against SARS-CoV-2 wild type and Delta challenges, resulting in a 100% survival rate. The mosaic 6RBD-np provides a potential platform for developing a pan-CoV vaccine against newly emerging SARS-CoV-2 variants and future CoV spillovers.

**Significance:** Despite the arsenal of COVID-19 vaccines, hospitalization and mortality associated with SARS-CoV-2 (acute respiratory syndrome coronavirus 2) variants remain high. There is an urgent need to develop next-generation COVID vaccines that provide broad protection against diseases by current and newly emerging SARS-CoV-2 variants. In this study, six receptor binding domains (RBDs) derived from α- and β-CoV were linked to proliferating cell nuclear antigen (PCNA) heterotrimeric scaffolds. They assemble to create a stable mosaic multivalent nanoparticle, 6RBD-np, displaying a ring-shaped disk with six protruding antigens. The prime-boost immunization in BALB/c and human angiotensin converting enzyme 2-transgenic mice with the 6RBD-np elicited strong, dose-dependent antibody responses and induced full-protection against both the SARS-CoV-2 wild type (WT) and Delta challenges. This study provides proof-of-concept that the mosaic 6RBD-np induces 100% protection against SARS-CoV-2 WT and Delta. It provides the potential of co-displaying heterologous antigens for novel vaccine designs, which can be deployed countering future pandemics.

## INTRODUCTION

The incidence of emerging infectious diseases has increased in the past 80 years^1^, and the majority of them are considered to be caused by a zoonotic spillover, posing ongoing human health threats^2, 3^. Recurrent spillovers of α- and β-coronaviruses (CoV), including severe acute respiratory syndrome (SARS)-CoV, Middle East respiratory syndrome (MERS)-CoV, SARS-CoV-2, and possibly human CoV (NL63, 229E, OC43, and HKU1), may have caused serious morbidity and mortality worldwide^4^. The ongoing pandemic of COVID-19 by SARS-CoV-2 has resulted in over 5.7 million deaths and 3.9 billion infections as of February 2022^5^. Despite high vaccine coverage, people suffer from an inevitable risk of antigenic drift variants, which poses additional threats to global health^6, 7^.

Upon infection, active SARS-CoV-2 spike glycoproteins, comprising of two subunits S1 and S2, play a key role in receptor binding and cell entry and are the primary target of neutralizing antibodies (nAbs)^8^. The S1 subunit consists of an N-terminal domain (NTD), a receptor binding domain (RBD), and S1 subdomains, SD1 and SD2. The RBD domain with an immunodominant receptor binding motif, a highly specific target of the nAbs, is observed in up and down conformations. It accounts for more than 90% of the neutralizing activity in sera from COVID-19 convalescent and vaccinated individuals^9–12^. Durable RBD-specific memory B cell and nAb responses are known to provide protection against infection^10, 13^. Notably, RBD monomer, tandem RBD dimer, and RBD trimer have been shown to induce potent Ab responses^14–16^. Self-assembling multivalent RBD-nanoparticles, mediated by a designed protein or scaffold protein conjugation using sortase A or spy-tag chemistry, further demonstrated extraordinarily high potentiation of vaccine-elicited immune responses^17–19^. These results highlight the importance of the RBD domain as a potential target as well as epitope density and organization.

Recent outbreaks of newly emerging variants indicated a high degree of waning vaccine-elicited immunity, contributing to increased infectiousness and immune evasion^20–22^. The new variants may be less vulnerable to the RBD-directed antibodies that exhibit high potency against ancestral SARS-CoV-2 strains^23–25^. Administration of a third dose of mRNA vaccines resulted in substantially lowered rates of confirmed cases and severe illness^19^. With the repeated booster vaccination, however, it has been suggested that immunological imprinting directed toward strain-specific antigens may lead to over-specialization of immunodominant B cells with limited breadth and a low activation of cross-reactive B cell with broad protection^26^. In this context, co-display of heterologous RBDs on nanoparticles has proven that the mosaic nanoparticles provide an avidity advantage to cross-reactive B cells, eliciting broader Ab responses than those induced by a mixture or cocktail of nanoparticles in animal experiments^26–28^. These innovative platforms prepared by designed scaffold, sortase A or Spy-tag-mediated fusion chemistry clearly induced high immunogenicity against RBDs and cross-reactive B cells with sufficient breadth. However, most of the platforms reported so far have significant drawbacks associated with employing a mixture of antigen-fused tags or scaffolds to assemble mosaic nanoparticles. In these cases, co-display of heterologous RBDs is challenging in mass production, due to the difficulty to achieve a uniform distribution of antigens on nanoparticles.

In this study, we embraced an approach that a novel self-assembling scaffold, proliferating cell nuclear antigen (PCNA), can form mosaic nanoparticles incorporating heterotypic antigens in uniform distribution. PCNA is a ring-shaped protein that encircles DNA as a processivity factor^29–31^. Most archaea encode a homotrimeric PCNA from bacteria to higher eukaryotes, including human, whereas a hyperthermophilic archaea, *Saccharolobus solfataricus*, possesses a heterotrimeric PCNA that is assembled by virtue of specific interactions between subunits PCNA1, PCNA2 and PCNA3. Six RBDs derived from α- and β-CoV that are considered to have emerged from the spillovers were selected and attached to PCNA, to form a stable mosaic multivalent nanoparticle, 6RBD-np. Prime-boost immunizations with 6RBD-np induced full-protection against SARS-CoV-2 wild type (WT) and Delta variant.

## RESULTS

### Design, assembly, and characterization of mosaic 6RBD-np antigen

To overcome the common disadvantages of mosaic nanoparticle platforms^25–28^, a heterotrimeric PCNA from *S. solfataricus* was chosen as a scaffold with three distinct subunits (PCNA1, 2, and 3). The subunits of ∼250 amino acids have similar structures, despite their low sequence similarity (8-22%), and are assembled in a sequential manner: PCNA1 and PCNA2 form a stable dimer, which recruits PCNA3 to form a heterotrimer, with dissociation constants in μM−nM ranges^29–32^. The antigens derived from the six CoV spike proteins were incorporated into the N- and C-terminus of PCNA1, PCNA2, and PCNA3 via SGG linker, which can confer multivalence to nanoparticles (see below).

Many nAbs directed against the RBD provide the highest neutralizing potencies, when targeting conserved epitopes exposed in the down conformation, and the presence of the SD1 stabilizes the RBD- down conformation^33–35^, which can also be critical for stability of spike proteins^36^. Six RBD-SD1s were thus selected, derived from α- and β-CoV that are considered to have originated from animals and caused zoonotic infections in humans^3, 4^: residues from 319 to 592 for SARS-CoV-2 WT and variant (VAR), 306 to 578 for SARS-CoV S, 367 to 657 for MERS-CoV S, 284 to 499 for hCoV 229E S, and 313 to 674 for hCoV HKU1 spike proteins (Fig. 1A). In addition, a variant RBD-SD1 of SARS-CoV-2 was included, where the K417N, L452R, T478K, E484K, and N501Y mutations were introduced using site-directed mutagenesis (Fig. 1A). Therefore, the fusion proteins, SARS-CoV-2 WT RBD-SD1-PCNA1-SARS-CoV RBD-SD1, MERS-CoV RBD-SD1-PCNA2-SARS-CoV-2 VAR RBD-SD1, and hCoV 229E RBD-SD1-PCNA3-hCoV HKU1 RBD-SD1, were constructed and termed S-PCNA1, M-PCNA2, and H-PCNA3, respectively (Fig. 1B).

**Fig. 1.**
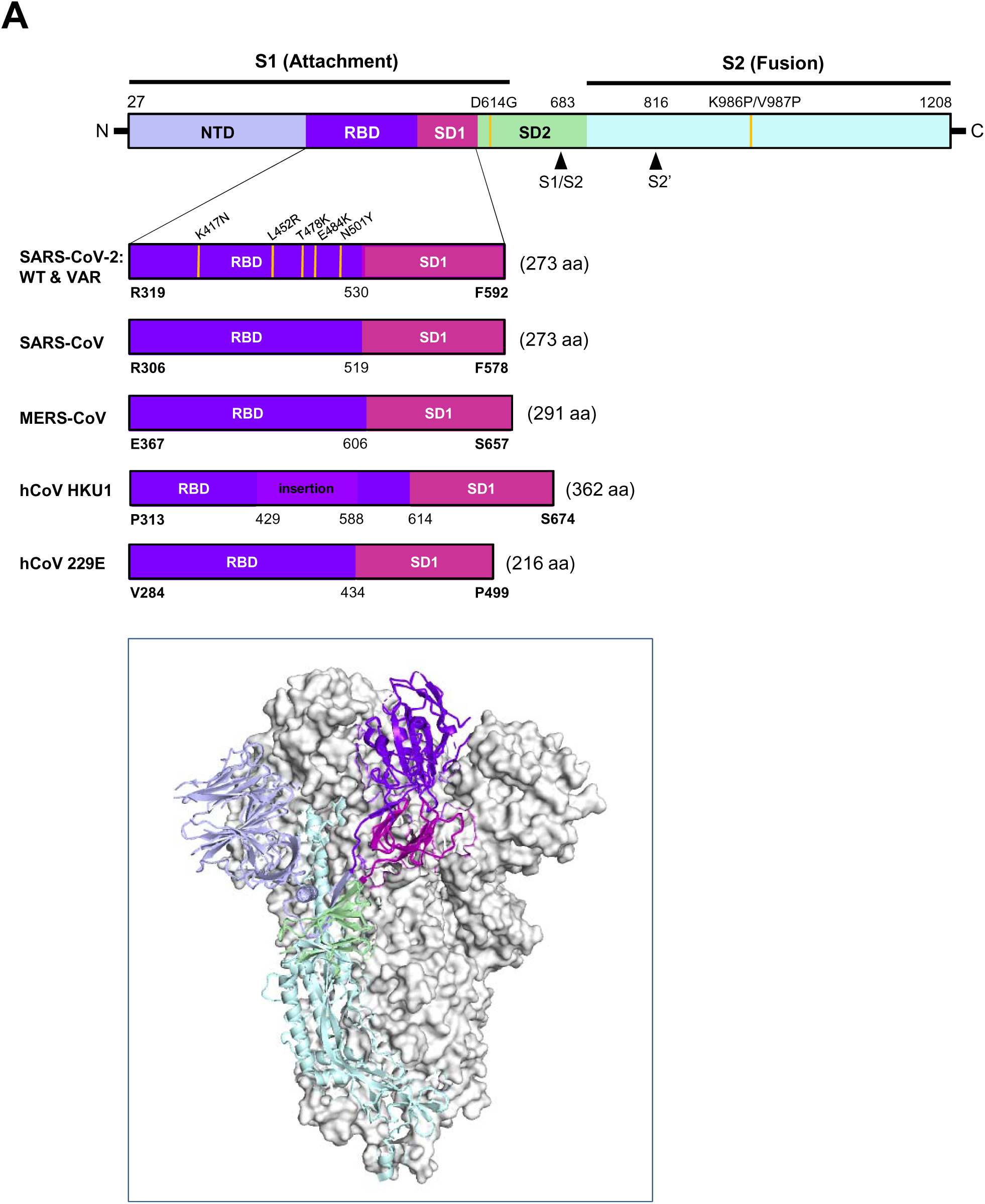
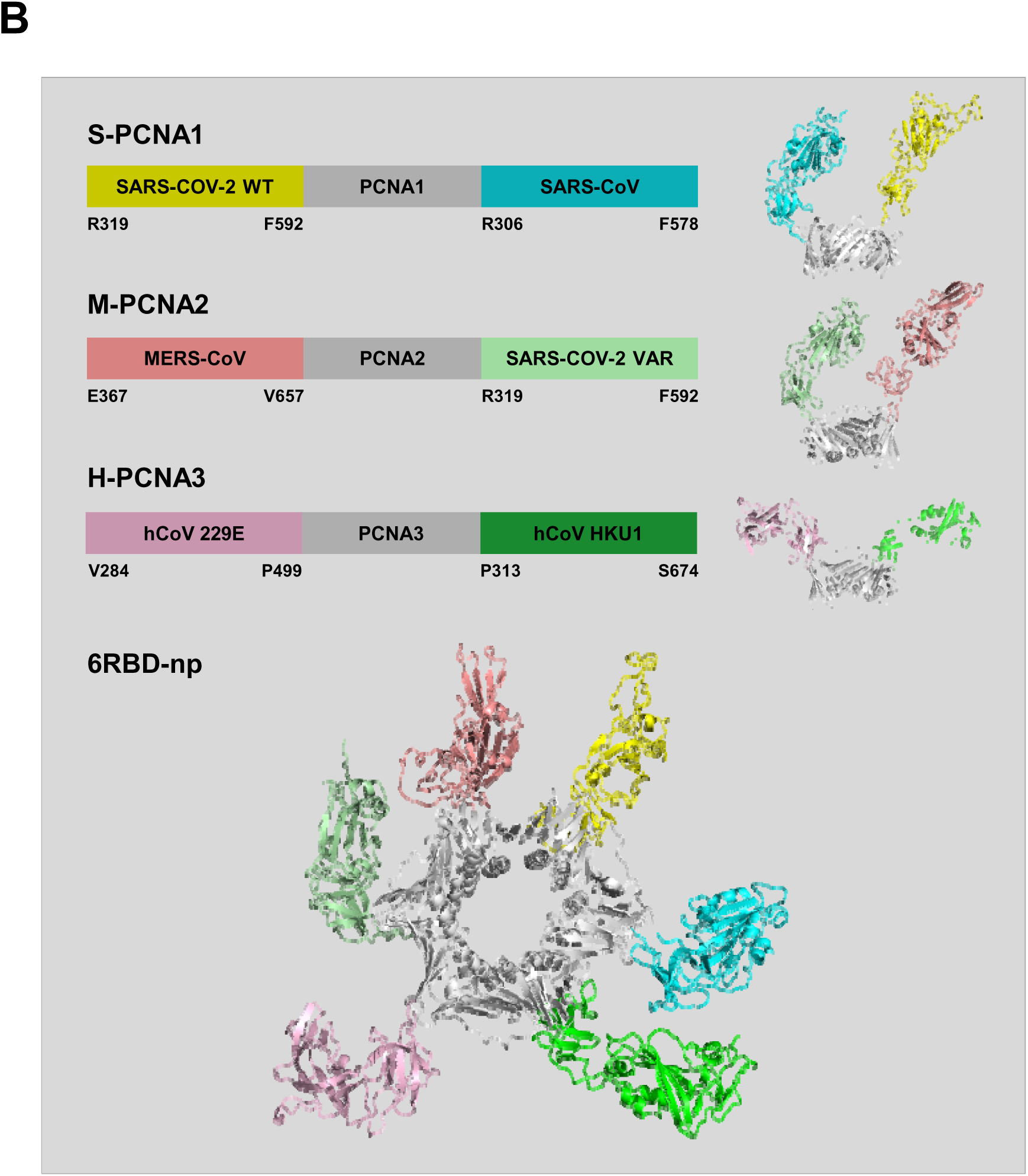
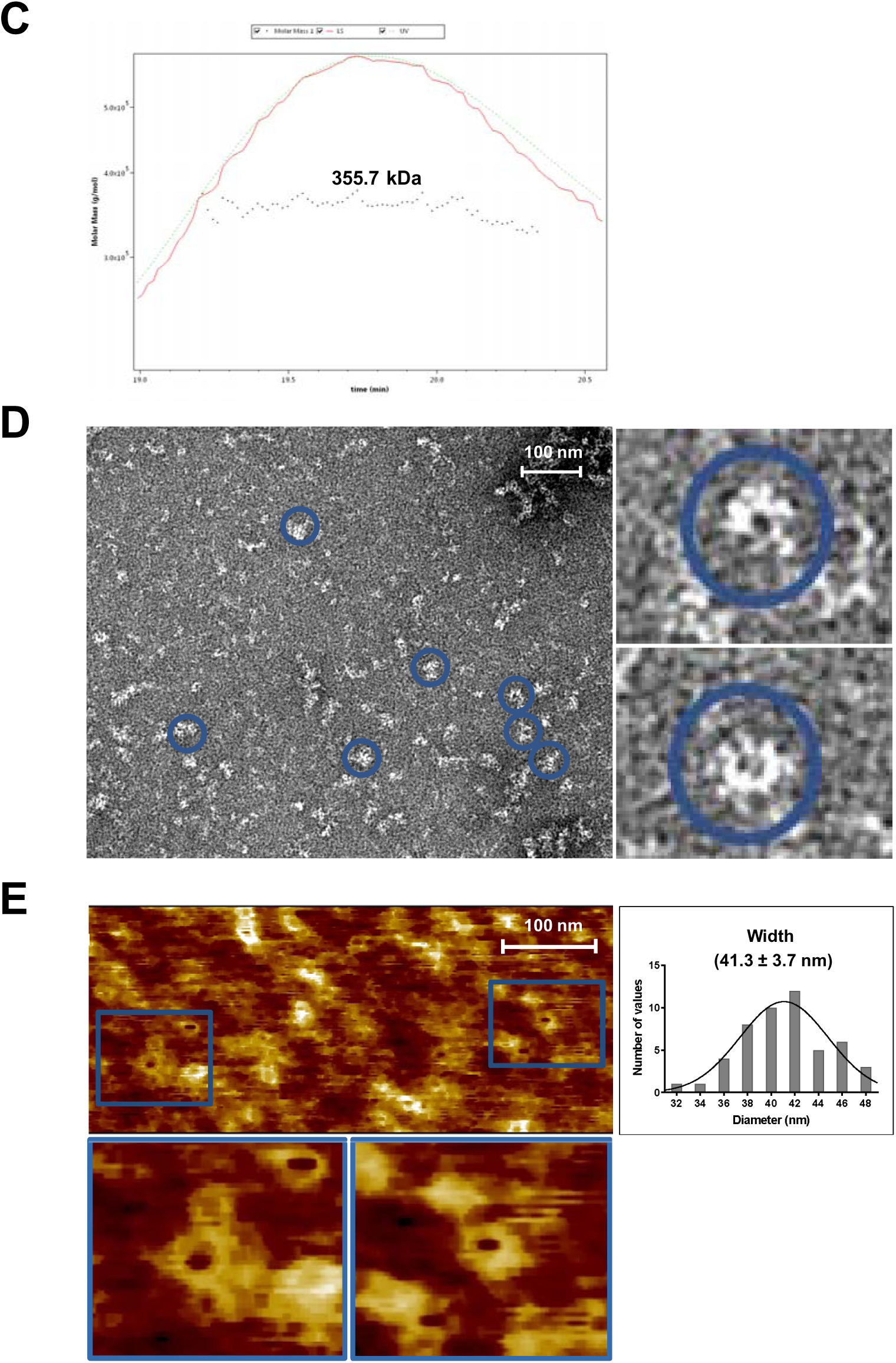
Design and characterization of 6RBD-np antigen. (A) Schematic of SARS-CoV-2 spike primary structure colored by domains (upper panel). S1 region contains NTD, N-terminal domain; RBD, receptor binding domain; SD1 and SD2, subdomain 1 and 2, and S2 region. The mutation sites for SARS-CoV-2 variants are indicated by yellow vertical lines. The RBD and SD1 regions are derived from from 319 to 592 for SARS-CoV-2 WT and variant, 306 to 578 for SARS-CoV, 367 to 657 for MERS-CoV, 313 to 674 for hCoV-HKU1, and 284 to 499 for hCoV-229E spike proteins. A prefusion trimeric structure of SARS-CoV-2 spike protein (lower panel). A single protomer is shown in ribbon representation and colored as in the primary structure diagram. The two remaining protomers are shown as molecular surface in gray. (B) Design of S-PCNA1, M-PCNA2 and H-PCNA3 showing schematic diagrams of CoV spike RBD-PCNA fusion proteins (upper panel): S-PCNA1, M-PCNA2 and H-PCNA3. S-PCNA1, SARS-CoV-2 RBD-SD1 WT-PCNA1-SARS-CoV RBD-SD1; M-PCNA2, MERS-CoV RBD-SD1-PCNA2-SARS-CoV-2 RBD-SD1 VAR; H-PCNA3, hCoV 229E RBD-SD1-PCNA3-hCoV HKU1 RBD-SD1. Composite molecular models of S-PCNA1, M-PCNA2 and H-PCNA3 are shown on the right. A molecular model of 6RBD-np displaying PCNA ring (gray) and 6 RBD-SD1 proteins colored as in the schematic diagram (lower panel). The models were generated from structures of PDB IDs: 6VXX, 5W9P, 5X5B, 6U7H, 5I08, and 2HIK for SARS-CoV-2, MERS-CoV, SARS-CoV, hCoV 229E, and hCoV HKU1 spike proteins, and PCNA, respectively. (C) Purified 6RBD-np was characterized using SEC-MALS to show its molecular weight of 355.7 kDa, suggesting a stable self-assembled 6RBD-np. (D) Negative staining electron (TEM) micrographs and (E) atomic force microscopic (AFM) images of the 6RBD-np with scale bars of 100 nm. They show a ring-shaped disk with six protruding RBD-SD1 antigens, like jewels in a crown, with an overall size of approximately 40 nm in width. The boxed rectangles on the right and below are magnified TEM and AFM images of 6RBD-np, respectively. The histogram of the width of 6RBD-np was prepared with 50-line profile data, based on AFM images.

The components for assembly, S-PCNA1, M-PCNA2, and H-PCNA3, were expressed in HEK 293F cells and purified using His-tag affinity, ion exchange and size exclusion chromatography (Extended Data Fig. 1A). They were mixed in a sequential order to generate mosaic multivalent nanoparticles, 6RBD-np (Extended Data Fig. 1B), showing the inter-subunit dissociation constants of S-PCNA1 and M-PCNA2 followed by H-PCNA3 for assembly of 1×10^-12^ and 1 x10^-7^, respectively (Extended Data Table 1). The purified 6RBD-np showed three well-resolved protein bands on SDS-PAGE and Western blot, suggesting distinct three specific components (Extended Data Fig. 2).

The purified antigens were characterized by SEC-MALS: S-PCNA1, M-PCNA2, H-PCNA3, and 6RBD-np showed molecular weights of 154.0 kDa (±12.2%), 182.2 kDa (±1.8%), 175.8 kDa (±1.6%), and 355.7 kDa (±3.6%), respectively (Fig. 1C and Extended Data Fig. 3). The 6RBD-np was further examined using atomic force microscopy (AFM) and transmission electron microscopy (TEM) to characterize its assembly, size, and morphology. A ring-shaped scaffold structure with protruding RBD-SD1 antigens was clearly observed, with an overall size of 40 nm (Figs. 1D and 1E).

### Antigenic characterization of mosaic 6RBD-np

The purified components, S-PCNA1, M-PCNA2 and H-PCNA3, were subjected to Western blot with anti-His-tag monoclonal Ab (mAb) (Abcam, Cambridge, UK) and anti-SARS-CoV-2 spike polyclonal Ab (pAb) (Sino Biological, China). All the antigens were detected by the anti-His-tag mAb and by the SARS-CoV-2 pAb, except for the H-PCNA3 (Fig. 2A). The H-PCNA3 was not detected by the anti-SARS-CoV-2 spike Ab, whereas the 6RBD-np and H-PCNA3 were identified by the anti-HKU1 spike mAb. The results clearly demonstrate that the engineered RBD antigens are present in the purified antigens, including the 6RBD-np. It is notable that the number of protein bands on Western blot detected by the anti-His-tag, anti-SARS-CoV-2, and anti-HKU1 Abs were 3, 2, and 1, respectively, strongly suggesting that the mosaic 6RBD-np indeed contain 6 RBD antigens in uniform distribution on nanoparticles.

**Fig. 2.**
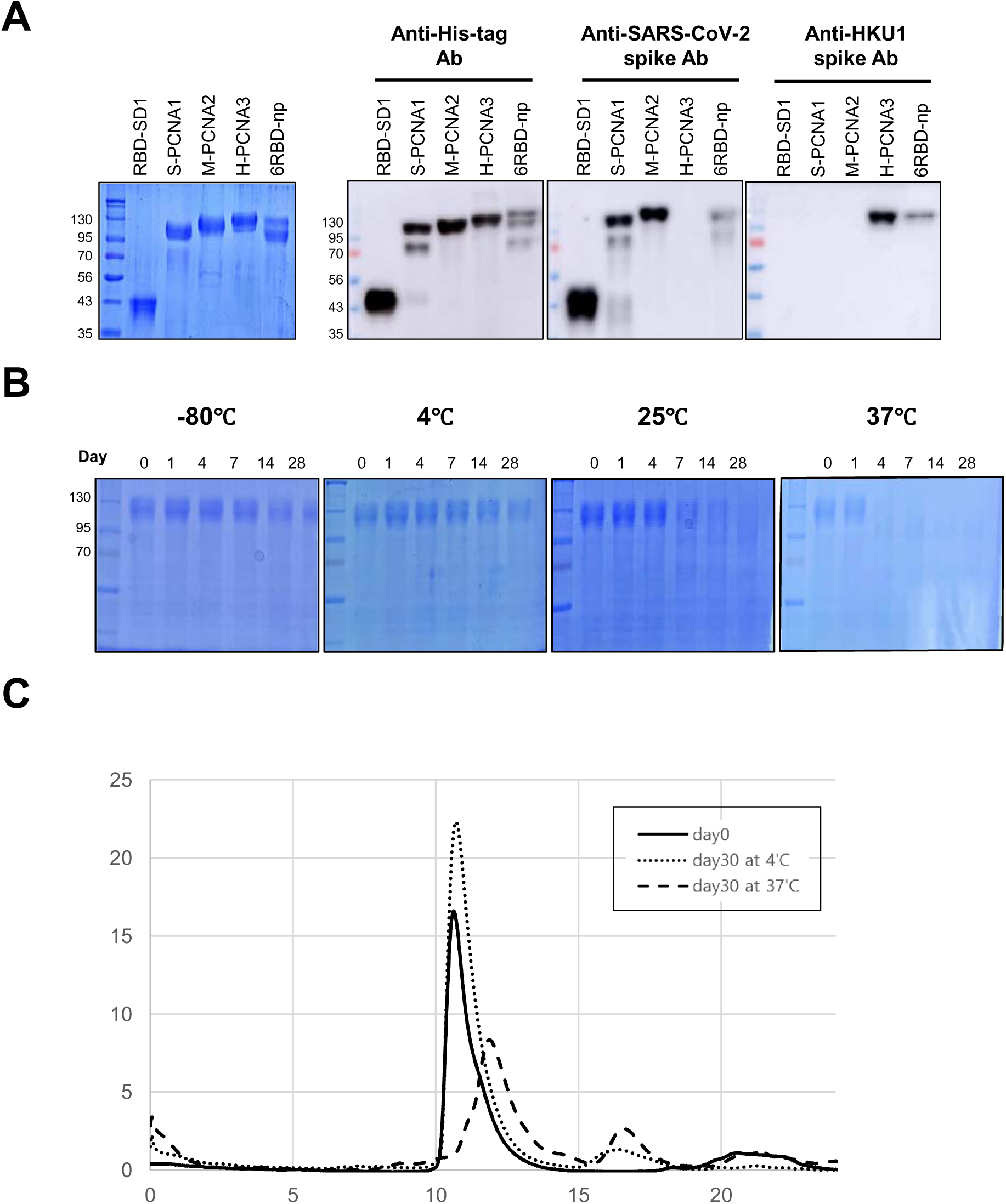
Antigenic characterization and stability of 6RBD-np. (A) SDS-PAGE and Western blot assay results of the purified components, RBD-SD1, S-PCNA1, M-PCNA2, H-PCNA3, and 6RBD-np, using anti-His-tag mAb, anti-SARS-CoV-2 spike pAb, and anti-MERS-CoV spike mAb. The S-PCNA1 and M-PCNA2 are detected by the SARS-CoV-2 pAb, while H-PCNA3 is not. Only the 6RBD-np and H-PCNA3 are detected by the anti-HKU1 spike mAb. (B & C) Time-course experiments of 6RBD-np monitored at −80°C, 4°C, 25°C, and 37°C over a 28-day storage period using SDS-PAGE and SEC. (B) The left lane, molecular weight markers, and following lanes monitored on days 0, 1, 4, 7, 14, and 28. (C) Elution profiles of 6RBD-np, obtained at day 0 (continuous line) and 30 at 4°C (dotted) and 37°C (dashed), after size exclusion chromatography on Superdex 200 increase 10/300 GL.

### Stability of mosaic 6RBD-np

The stability of the component proteins and 6RBD-np was monitored at −80°C, 4°C, 25°C, and 37°C over a 28-day storage period, using SDS-PAGE and SEC. They were stable and resistant to degradation at −80°C over 28 days (Fig. 2B and Extended Data Fig. 4). Incubation at 4°C also showed excellent stability, and partial degradation started at 21 or 28 days in S-PCNA1 only. In contrast, severe degradation was observed at 25°C after 4 days of incubation. Significant degradation started in all the antigens at 37°C after 1 day of incubation. The stability for these antigens was also examined using SEC after 30 days (Fig. 2C and Extended Data Fig. 5). The 6RBD-np and H-PCNA3 were found to be stable for more than 30 days at 4°C, at which other components S-PCNA1 and M-PCNA2 are less stable. Therefore, the results suggest that the mosaic nanoparticle, 6RBD-np, is likely to be equally or more stable upon assembly than the components, representing a stable mosaic multivalent nanoparticle.

### *In vivo* immune response against antigens

Immunogenicity of the antigens was evaluated in BALB/c mice who received intramuscular prime-boost immunizations with a 3-week interval. Blood samples were collected at three time points: two weeks post-prime and two and four weeks post-boost (Fig. 3A). The mice were divided into six groups of RBD-SD1 of SARS-CoV-2, spike ectodomain with two proline substitutions (S-2P) of SARS-CoV-2, S-PCNA1, and 6RBD-np (n=10), including the naïve and PCNA control (n=5) groups. In RBD-based ELISA assays, both the S-PCNA1 and 6RBD-np groups showed the highest Ab titers after the boost, followed by the RBD-SD1 and S-2P groups (Fig. 3A, middle panel). The S-PCNA1 group, containing the SARS-CoV-2 and SARS-CoV RBD-SD1 immunogens, is considered to receive half the amount of the SARS-CoV-2 RBD-SD1, compared to the RBD-SD1 group. Likewise, the 6RBD- np mice are estimated to receive a sixth of the amount of the SARS-CoV-2 RBD-SD1. In this context, the results strongly suggest that both the 6RBD-np and S-PCNA1 groups showed significant and robust Ab responses, compared with the others. When the Ab responses in the post-boost sera were further compared with respect to dosage (15 μg vs 5 μg) and adjuvants (SAS, AddaVax and CpG-ODN), the 15 μg (3×) 6RBD-np group achieved the highest Ab response (Fig. 3A, lower panel). The adjuvants in this study enhanced the immune responses, but no significant difference was observed among the adjuvant groups. Therefore, our results demonstrate that the 6RBD-np induces strong antigen-specific Ab responses in mice by prime-boost immunizations.

**Fig. 3.**
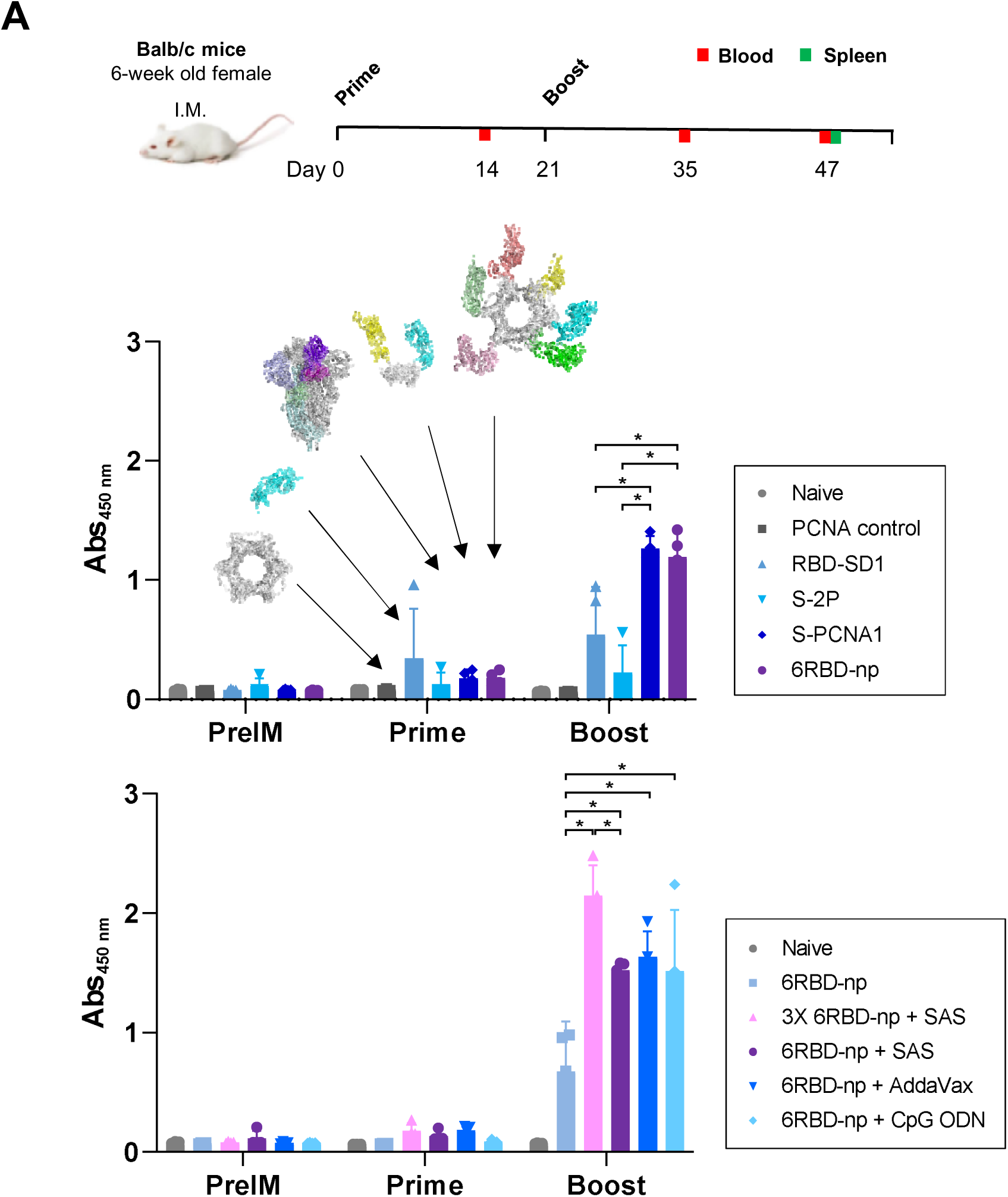
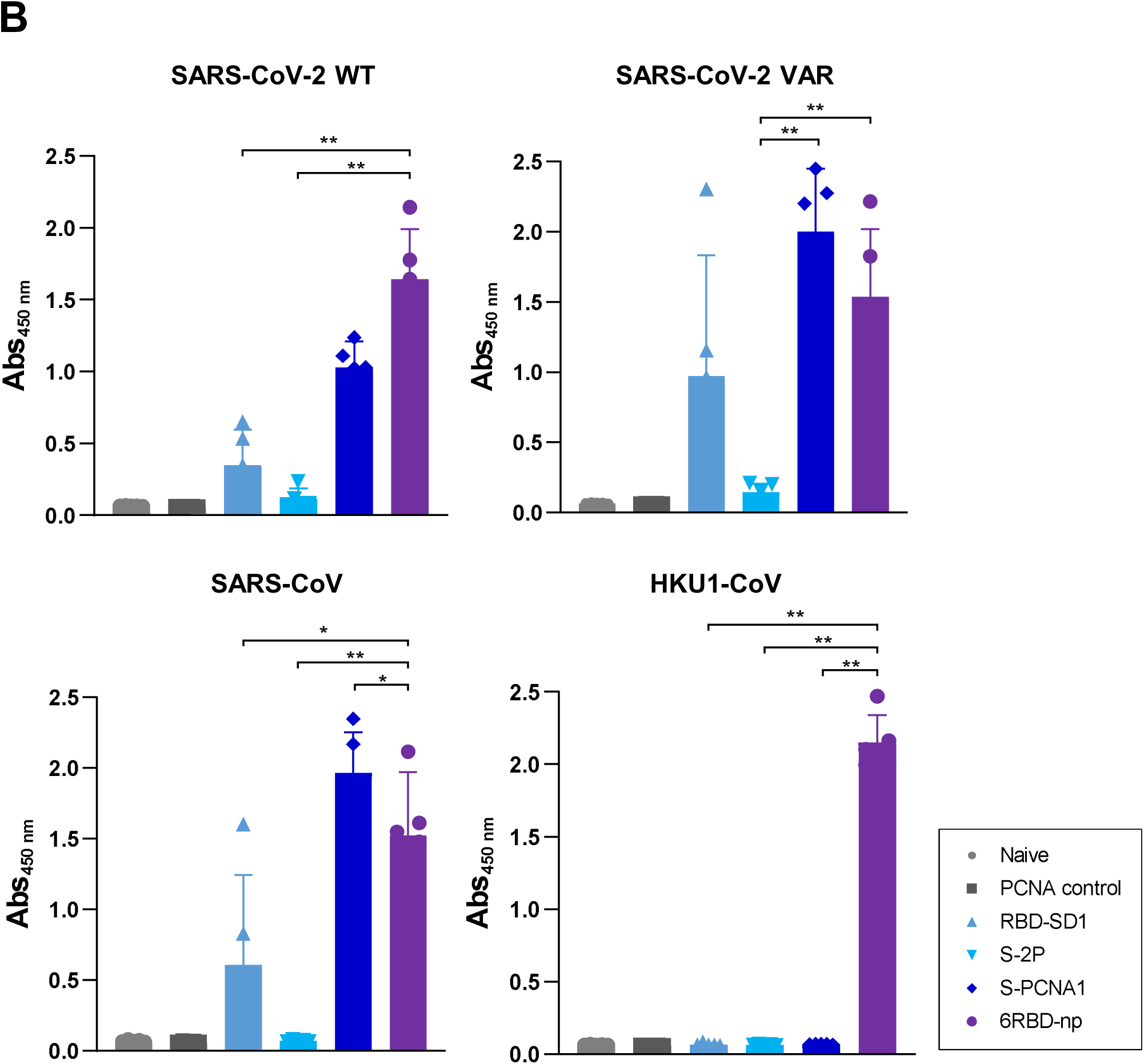
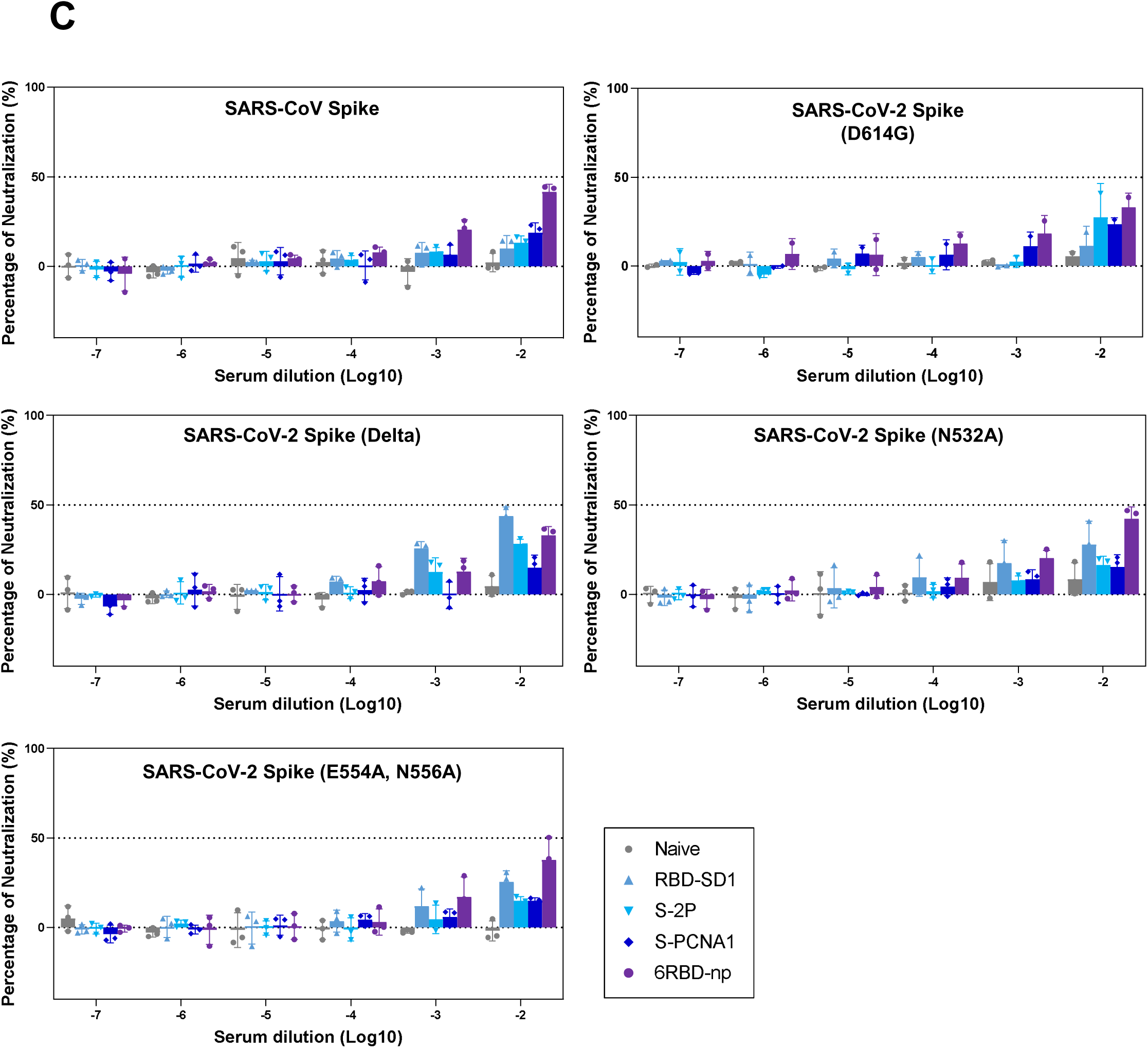
Immunogenicity of antigens in BALB/c mice. (A) Prime-boost immunization study design with BALB/c mice (upper panel). The mice were immunized in six groups. Blood was collected two weeks after the prime and boost immunizations and 26 days after the boost. Post-prime and post-boost anti-RBD Ab titers in BALB/c mice, for different immunogens of the groups of the naïve and PCNA controls, RBD-SD1 of SARS-CoV-2, S-2P of SARS-CoV-2, S-PCNA1, and 6RBD-np, represented as molecular structures of the respective immunogens with arrows (middle panel). Additional experiments were performed for different doses and adjuvants (lower panel), in groups of 5 μg and 15 μg of 6RBD-np, 5 μg of 6RBD-np in the presence of SAS, AddaVax, and CpG-ODN. The 15 μg 6RBD-np group shows the highest Ab response, compared to other groups. (B) Post-boost anti-RBD Ab titers against the purified RBD-SD1s of the SARS-CoV-2 (WT and VAR), SARS-CoV, and hCoV HKU1. Mice sera from the S-PCNA1 and 6RBD-np groups show high Ab titers against the RBD-SD1s of the SARS-CoV-2 (WT and VAR) and SARS-CoV, whereas only the 6RBD-np group reveals high titers against the HKU1 RBD-SD1. (C) Pseudovirus-based neutralization activities. Pseudoviruses expressing SARS-CoV-2 spike WT, Delta, and SD1 mutants were inoculated with different sera from the naïve and PCNA controls, RBD-SD1 and S-2P of SARS-CoV-2, S-PCNA1, and 6RBD-np groups, which were serially diluted prior to transduction on HEK293T-hACE2 cells. Neutralization was performed twice and a representative is shown. In (A) and (B), data were compared using the Mann-Whitney U test (*p < 0.05; **p < 0.01) and median values are represented by a solid horizontal bar. Statistically significant differences were indicated only among different immunogen groups.

We then assessed antigen-specific Ab titers in the mice sera, using purified recombinant RBD-SD1s of SARS-CoV-2 (WT and VAR), SARS-CoV, and hCoV HKU1. The groups of S-PCNA1 and 6RBD-np showed distinctly high Ab titers against SARS-CoV-2 (WT and VAR) and SARS-CoV RBD-SD1s, whereas only the 6RBD-np group displayed high Ab titers against the HKU1 RBD-SD1 (Fig. 3B). The results also suggest that the 6RBD-np displays mosaic heterologous RBD antigens. Further, neutralizing activities of the mice sera against pseudoviruses encoding spike of SARS-CoV and spike mutants of SARS-CoV-2 including D614G (Wuhan variant), L452R/T478K/D614G (Delta variant), N532A, and E554A/N556A (SD1 variants) were determined. The results show that the 6RBD- np group neutralized the entry of pseudoviruses, expressing SARS-CoV spike or SARS-CoV-2 spikes containing D614G, Delta or SD1 mutations, with high efficacy (Fig. 3C). The neutralization of the 6RBD-np group was lower than the RBD-SD1 group against the Delta spike. Nevertheless, our data strongly suggest that the nAbs in the 6RBD-np group neutralized cell entry of pseudoviruses, irrespective of RBD and SD1 mutations and irrespective of SARS-CoV and SARS-CoV-2 spikes used in this study.

### 6RBD-np completely protects mice against SARS-CoV-2 WT and Delta challenges

To evaluate the immunogenicity and protective efficacy of the antigens against the SARS-CoV-2 WT and Delta challenges, hACE2 knock-in mice were given with intramuscular prime-boost immunizations with a 3-week interval (Fig. 4A): groups of RBD-SD1 of SARS-CoV-2, S-2P of SARS-CoV-2, S-PCNA1, and 6RBD-np (n=10 each for the WT and Delta variant), including the naïve (n=6) and virus control (n=8) groups. At 42 days post-prime, all groups except for the naïve group were challenged with intranasal inoculation of 10 MLD_50_ of the SARS-CoV-2 WT or Delta. The S-2P, S-PCNA1 and 6RBD-np groups exhibited survival rates of 100% against the WT challenge, whereas the RBD-SD1 and 6RBD-np groups survived at the same rates against the Delta challenge (Fig. 4A, middle and lower panels). Notably, no significant changes in body weights were observed in the 6RBD-np group, the only group showing survival rates of 100%, irrespective of the WT or Delta challenge. The S-2P and S-PCNA1 groups exhibited average survival rates of 86% and 71% against Delta, respectively, while the RBD-SD1 group showed a survival rate of 57% against the WT challenge, exhibiting a relatively large body weight loss. Mice in the virus control group inoculated intranasally with the WT or Delta variant challenge started to lose weight at 2 or 3 days post-inoculation (dpi), respectively, showing rapid body weight loss and decline in survival rates.

**Fig. 4.**
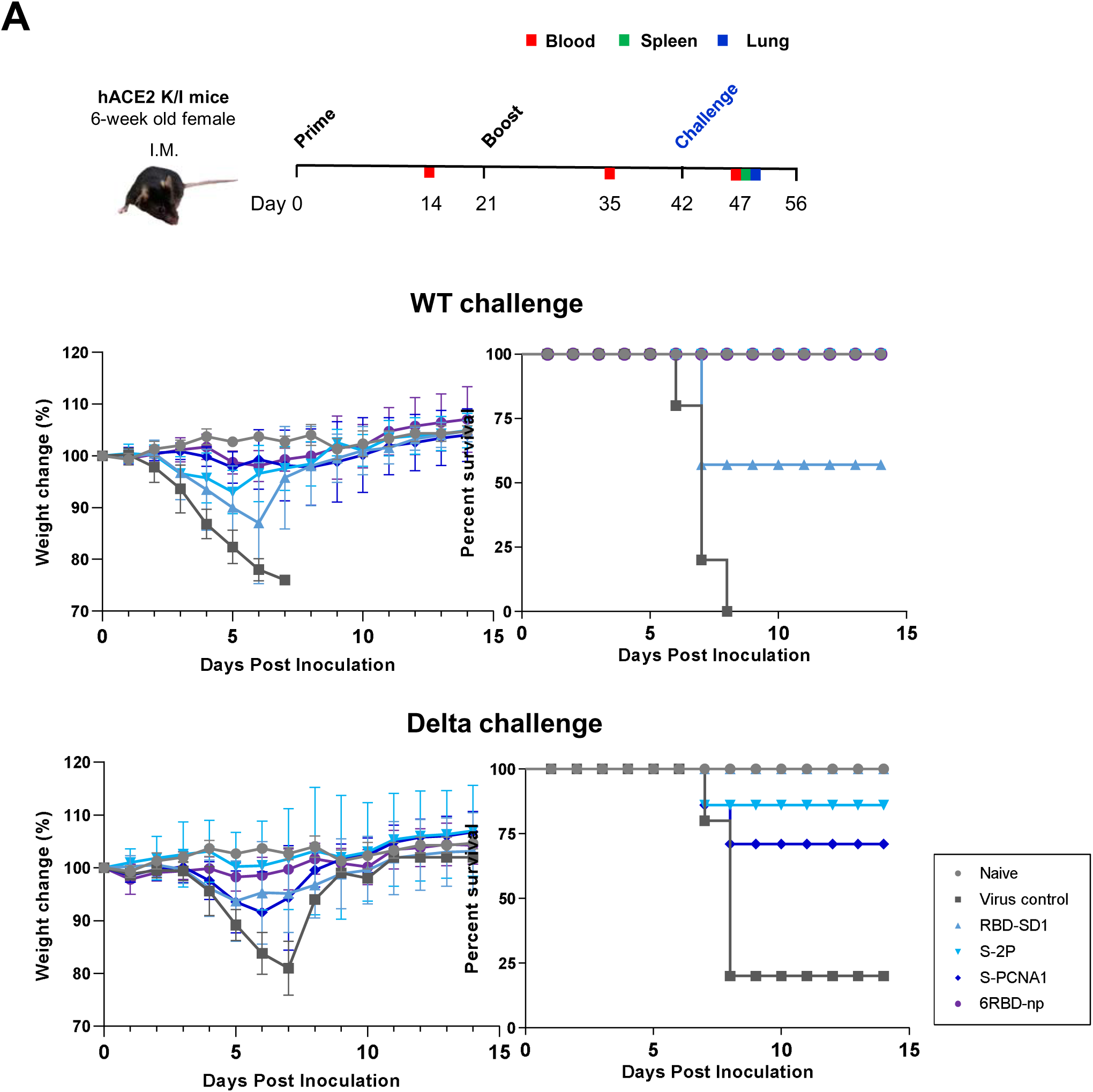
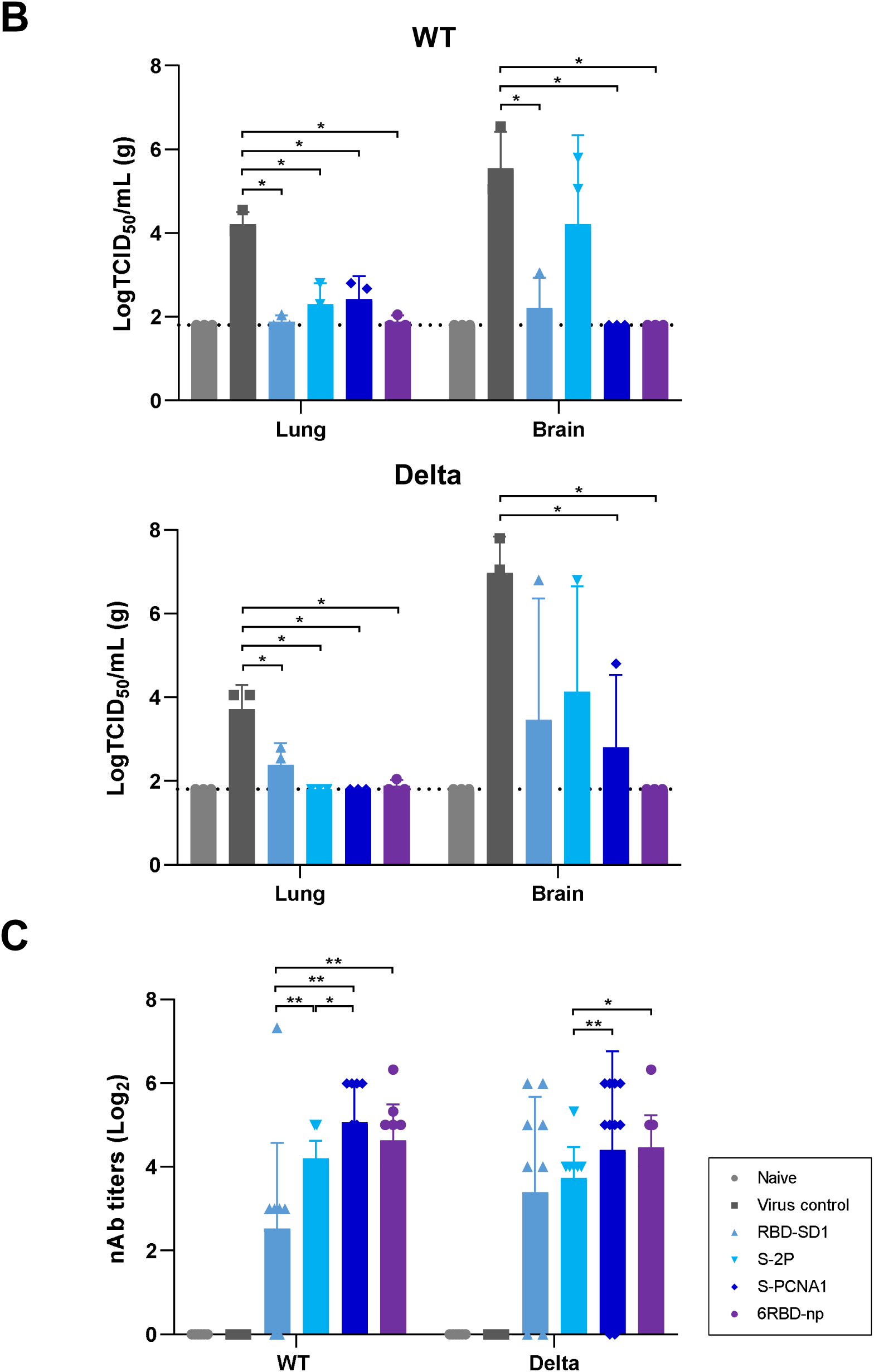
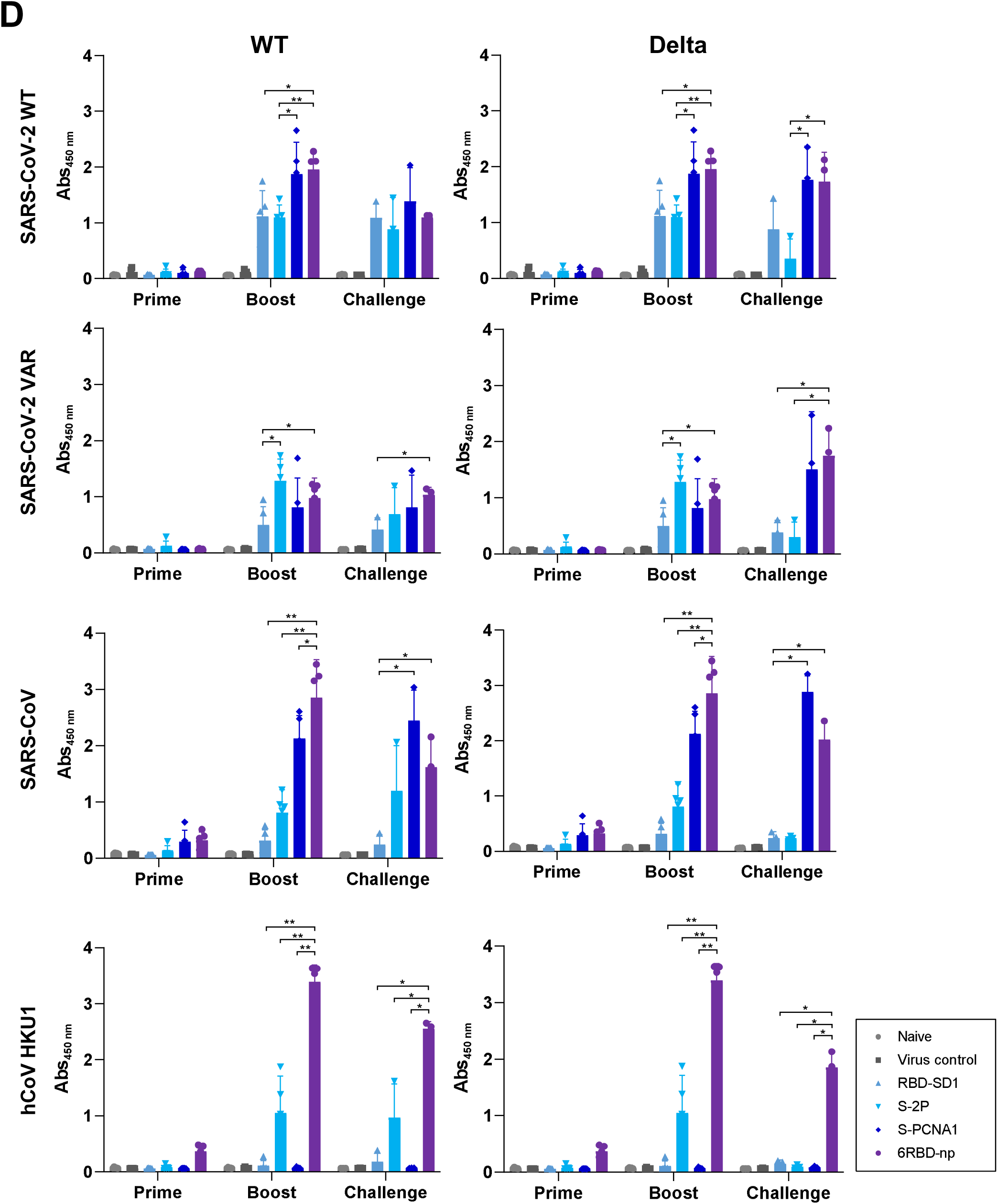
Protective efficacy and immunogenicity of antigens against SARS-CoV-2 WT and Delta challenges in hACE2 transgenic mice. (A) Prime-boost immunization and challenge study design with hACE2 transgenic mice (upper panel). Weight loss and survival of hACE2 transgenic mice, following the SARS-CoV-2 WT or Delta challenge up to 14 days post-infection, in groups of RBD-SD1 and S-2P of SARS-CoV-2, S-PCNA1, and 6RBD-np (n=10), including the naïve (n=6) and virus-infected control (n=8) groups. The mosaic 6RBD-np group shown in violet demonstrates little changes in body weight and survival, the only group showing survival rates of 100%, regardless of the virus strains (middle and lower panels). The naïve mice are shown by a blue line. (B) Viral titers in mice lungs and brain, collected from 3 mice per group, obtained 5 days post-infection (dpi), following the SARS-CoV-2 WT or Delta challenge with 10 MLD_50_, expressed in logTCID_50_/mL. The 6RBD-np group reduces the virus titers very similar to those in the naïve group, in the WT and Delta challenges. (C) Neutralizing Ab titers in mice sera collected at 35 days post-prime against the WT and Delta viruses. The S-PCNA1 and 6RBD-np groups show significantly higher immunogenicity than the RBD-SD1 and S-2P groups against the WT and delta viruses, respectively. (D) Anti-RBD Ab titers in mice sera against the WT and Delta viruses after prime, boost and challenges, using ELISA assays. The antigens used were the purified RBDs of SARS-CoV-2 (WT and VAR), SARS-CoV, and hCoV HKU1. The groups of S-PCNA1 and 6RBD-np show high titers. In (B)−(D), the data were compared using the Mann-Whitney U test (*p < 0.05; **p < 0.01) and median values are indicated by a bar. In (C) and (D), statistically significant differences were indicated only among different immunogen groups.

When the viral titers were measured in the lung and brain tissues collected at 5 dpi, the lowest titers were observed in the 6RBD-np and naïve groups, irrespective of the WT and Delta variant challenges (Fig. 4B). The viral titers in the harvested tissues were partially inhibited in the RBD-SD1, S-2P, and S-PCNA1 groups, compared to those in the control group. The mice in the immunized groups exhibited less severe histopathological lesions of interstitial pneumonia and lower N proteins, detected by immunohistochemistry (IHC), than those in the virus challenged group (Extended Data Fig. 6). In particular, the mice immunized with 6RBD-np revealed little histopathological damage and lack of N proteins in the lung, compared to other immunized groups.

Serum nAb titers are currently accepted as a correlate of protection against SARS-CoV-2 variants^12, 37, 38^. We next assessed nAb titers in post-boost sera, using the WT and Delta viruses. The S-PCNA1 and 6RBD-np groups revealed substantially high neutralizing potency and breadth against the SARS-CoV-2 WT as well as Delta variant (Fig. 4C). The S-PCNA1 and 6RBD-np groups could reach maximum titers of 64−80 neutralizing units against both WT and Delta viruses. Overall, the immunogenicity of the 6RBD-np in mice sera was significantly high both in the authentic SARS-CoV-2-based and pseudovirus-based neutralization assays, as shown in Fig. 3C. Moreover, the antigen-specific Ab titers were examined using the SARS-CoV-2 (WT and VAR), SARS-CoV, and hCoV HKU1 RBDs. Although the S-2P group showed high Ab titers specific to SARS-CoV-2 VAR at 2 weeks post-boost, the S-PCNA1 and 6RBD-np immunogens induced high Ab titers against SARS-CoV-2 and SARS-CoV RBDs at 5 days post-challenge (Fig. 4D). As expected, the 6RBD-np induced the highest titers against the HKU1 RBD. Taken together, the results demonstrate that the 6RBD-np antigen elicited significantly high titers of nAbs, despite being given a sixth dose of the RBD-SD1, which confer complete protection against both the SARS-CoV-2 WT and Delta in the hACE2 transgenic mouse model.

## Discussion

An arsenal of highly effective COVID-19 vaccines provides a robust threshold of protection against SARS-CoV-2^39–42^. However, multiple variants, including Delta (B.1.617.2) and Omicron (B.1.1.529), have been identified globally^23–25, 43^, and the Omicron variant, harboring 37 amino acid changes to the spike protein, has substantially increased ability to evade immunity from vaccination or prior infection^43–45^. As the current vaccines are mostly directed toward the viral spike protein, specifically in receptor interaction, the variants have raised a global concern over waning immunity and a global threat to public health^38^. Nevertheless, nAbs against multiple variants are currently accepted as a correlate of protection^12, 37^, and the RBD indeed accounts for more than 90% of the neutralizing activity in sera from COVID-19 convalescent and vaccinated individuals^9–12^. RBD monomer, dimer and trimer antigens elicited potent Ab responses^9, 10, 13^, while highly ordered and repetitive structures of RBDs mimicked the organization and density of antigens on natural virions, to be of great advantage in promoting protective immunity^17, 46, 47^. Self-assembling multivalent RBD or spike nanoparticles using SpyTag/SpyCatcher or sortase-mediated ligation and computationally designed scaffolds, ferritin, or lumazine synthase fusion platforms have shown high potentiation of immune responses against COVID-19^17–19, 47, 48^. However, administration of homotypic multivalent antigens or possibly multiple booster doses with strain-specific antigens can lead to over-specialization of the B cells with limited breadth of antigen recognition^26^. In this context, it is notable that mosaic nanoparticles with heterologous RBDs provide an avidity advantage to elicit cross-reactive B cells with broader Ab responses than those induced by an admixture of nanoparticles^27, 28^. These RBD nanoparticles were assembled by mixing 20 trimeric and 12 pentameric designed fusion proteins or by mixing 4 or 8 heterologous β-CoV RBDs fused to scaffolds, which elicited Abs with superior cross-reactive recognition of heterologous RBDs relative to homotypic or cocktail RBD nanoparticles.

Molecular self-assembly of current mosaic nanoparticles is commonly achieved by mixing heterologous antigen-fused tags or scaffolds, However, a formulation of mosaic nanoparticles with a uniform distribution of heterotypic antigens cannot be ensured, and quality assurance in mass production will be more challenging in human trials. In this study, six heterologous RBD-SD1s are fused to the N- and C-termini of PCNA1, 2, and 3, which are assembled in a defined order to form a nanoparticle, driven by specific interactions of the heterotrimeric subunits^29, 30^. Each PCNA subunit confers two heterologous RBD-SD1 antigens at the N- and C-termini, and the assembled heterotrimer can present up to six heterologous RBD-SD1s. The 6RBD-np characterized by SEC-MALS, TEM and AFM has a molecular weight of 355 kDa with an overall size of 40 nm, which appears to be a ring-shaped disk with six protruding RBD-SD1s, like jewels in a crown (Fig. 1D & 1E). Antigenic characterization by Western blot using His-tag mAb, SARS-CoV-2 pAb, and HKU1 Ab reveals that six heterologous RBD antigens are uniformly distributed on the 6RBD-np nanoparticle.

Efficient cellular uptake, entry into, and retention at lymph nodes of antigens are crucial for improved immunogenicity and protection, which is influenced by the size of antigen nanoparticle platforms^48, 49^. Self-assembled protein nanoparticles are defined as a material that is deliberately manipulated to have dimensions between 1 to 100 nm or up to 1,000 nm if they exhibit physical, chemical, or biological effects^50^. The repetitive array of antigens on nanoparticles facilitates efficient binding and activation of multiple B cell receptors, highlighting the importance of antigen density in memory immune responses^49, 51^. Critical factors for successful use of nanoparticle vaccines are thus size, antigen density, and surface properties. Although discrepancies may arise, the parameters for the ideal nanoparticle platform would be 20−50 nm in diameter, with antigens spaced 30 nm apart^48^. The 6RBD-np with the size of ∼40 nm is shown to have inter-antigen spacing of 20−30 nm and to uniformly present up to six heterologous antigens.

In the animal experiments using BALB/c mice, the 6RBD-np group demonstrated high Ab titers for each given antigen, compared to other groups after the boost immunization (Fig. 3A & 3B). In the dose-response experiment, the 15 μg dose in BALB/c mice achieved the highest Ab response. Given that the 6RBD-np mice receive a sixth of the amount of the SARS-CoV-2 RBD-SD1, compared to the RBD-SD1 group, it clearly demonstrates that the 6RBD-np induced significant Ab responses. Importantly, in hACE2 transgenic mice, the prime-boost immunization of the 6RBD-np induced protective immunity against both the SARS-CoV-2 WT and Delta challenges (Fig. 4A). The virus titers at 5 dpi in the lungs and brain significantly dropped to the baseline in the 6RBD-np group, essentially identical to that in the naïve group. The 6RBD-np elicited significantly high neutralizing potency in mice after the boost, when using the authentic SARS-CoV-2-based assays, which was consistent with that in the pseudovirus-based neutralization assays. Therefore, the 6RBD-np induced significant and robust Ab responses in BALB/c mice as well as in hACE2 transgenic mice, demonstrating the only group with 100% survival rates among the antigen groups challenged with the SARS-COV-2 WT or Delta variant. Our results are in good agreement with recent studies showing that mosaic nanoparticles led to significantly higher immune responses and protection efficiency^27, 28^. This mosaic array of 6RBD-np would provide a potential route to produce pan-CoV vaccines to combat future spillovers.

## Materials and methods

### Cell lines and animals

Expi293F cells derived from the 293F cell line (Life Technologies, Carlsbad, CA, USA) were grown in Expi293 expression medium (Life Technologies), cultured at 37°C with 8% CO_2_ and shaking at 150 rpm. Human embryonic kidney cells (HEK293T) is a female human embryonic kidney cell line (ATCC) and the HEK-ACE2 (derived from HEK293T cells) adherent cell line was obtained through BEI Resources, NIAID, NIH: HEK293T expressing human angiotensin-converting enzyme 2 (hACE2), HEK293T-hACE2 cell line (NR-52511).

HEK293T and HEK-ACE2 cells were cultured in Dulbecco’s modified Eagle’s medium (DMEM) (Gibco, Texas, USA) supplemented with 10% fetal bovine serum (FBS) (Atlas Biologicals, Colorado, USA) and 1% penicillin-streptomycin (P/S) (Hyclone, Waltham, MA, USA). To generate HEK293T cell line stably expressing hACE2, HEK293T cells were transfected with pCEP4-myc-hACE2 and cells were selected using 100 μg/mL of hygromycin B (Invitrogen, Logan, Utah, USA). The detailed procedure was delineated previously^52^. Vero E6 (ATCC, Manassas, VA, USA) is an African green monkey epithelial kidney cell line. Vero E6 cells were maintained in DMEM containing 10% FBS, 4.0 mM L-glutamine, 110 mg/L sodium pyruvate, 4.5 g/L D-glucose and 1% antibiotic-antimycotic at 37°C and 5% CO_2_. Six-week old female B6.Cg-Tg(K18-ACE2)2Prlmn/J mice were obtained from Jackson Laboratory (Bar Harbor, Maine, USA).

### Plasmid construction

The S-PCNA1, M-PCNA2, and H-PCNA3 were produced using polymerase chain reaction (PCR). For 6RBD-SD1 proteins, residues from 319 to 592 for SARS-CoV-2 WT and variant (VAR), 306 to 578 for SARS-CoV S, 367 to 657 for MERS-CoV S, 284 to 499 for hCoV 229E S, and 313 to 674 for hCoV HKU1 spike proteins as well as PCNA1, PCNA2, PCNA3_S170V were amplified from the pSecTag2A vector (Addgene) using the primers described in Extended Data Table 2. Fusion proteins, SARS-CoV-2 RBD-SD1 WT-PCNA1-SARS-CoV RBD-SD1, MERS-CoV RBD-SD1-PCNA2-SARS-CoV-2 RBD-SD1 VAR, and hCoV 229E RBD-SD1-PCNA3-hCoV HKU1 RBD-SD1, were constructed with SGG linker sequences by overlapping PCR, which were termed S-PCNA1, M-PCNA2, and H-PCNA3, respectively. They were ligated to pSecTag2A vectors using restriction enzymes, BamH1 and Not1, with myc and 6× histidine tag at the downstream of each fusion protein. Additionally, the RBD-SD1 of SARS-CoV-2 VAR was included, where the K417N, L452R, T478K, E484K, and N501Y mutations were introduced using site-directed mutagenesis. Plasmids encoding S-PCNA1, M-PCNA2, and H-PCNA3 were amplified in *Escherichia coli* strain DH5α, which were prepared in high purity and concentration using the PureLink HiPure Plasmid Kit (Invitrogen). All sequences were confirmed by automated sequencing (Macrogen, Seoul, Korea).

### Transfection, expression, and purification of 6RBD-np

Expi293F cells were transfected with the purified plasmids using ExpiFectamine 293 transfection kit (Invitrogen), where the fusion proteins were expressed and harvested at 96 hrs post-transfection under 8% CO_2_ at 37°C. They were centrifuged at 4,000 rpm for 1 hr to get the supernatant containing each fusion protein, S-PCNA1, M-PCNA2, or H-PCNA3, which was applied to a Ni-NTA affinity chromatography column (Qiagen, Hilden, Germany), equilibrated with 20 mM Tris-HCl (pH 8.0) and 100 mM NaCl. The column was washed with the equilibration buffer containing 20 mM imidazole and the fusion protein was eluted with elution buffer, 20 mM Tris-HCl (pH 8.0), 100 mM NaCl, and 400 mM imidazole. The eluted protein was dialyzed against 20 mM Tris-HCl (pH 8.0) and 10 mM NaCl for 16 hrs at 4°C with stirring. Further purification was performed by ion-exchange chromatography using mono Q 5/50 GL (GE HealthCare, Uppsala, Sweden) for each fusion protein in 20 mM Tris-HCl (pH 8.0) and 1 M NaCl, and by size exclusion chromatography using Superdex 200 Increase 10/300 GL column (GE Healthcare, Uppsala, Sweden) in Dulbecco′s phosphate buffered saline (DPBS), connected to an Ä KTA FPLC system (GE HealthCare). The purified S-PCNA1 and M-PCNA2 were mixed with a 1.2:1 molar ratio at 4°C for 6 hrs, to which H-PCNA3 was added in 1.2:1 ratio at 4°C for 6 hrs. The assembled 6RBD-np was purified by size exclusion chromatography using Superdex 200 Increase 10/300 GL column (GE Healthcare) in DPBS. Sodium dodecyl sulphate-polyacrylamide gel electrophoresis (SDS-PAGE) was used to assess protein purity.

### Cloning, expression, and purification of RBD-SD1 antigens

The constructs for RBD-SD1 of SARS-CoV-2 WT, SARS-CoV-2 VAR, SARS-CoV, and hCoV HKU1 as well as SD1s of SARS-CoV-2 WT, SARS-CoV, MERS-CoV, and hCoV HKU1 were produced by PCR using RBD-SD1-PCNA as a template. Plasmids encoding each antigen were amplified in *E. coli* strain DH5α, which were prepared in high purity and concentration using the PureLink HiPure plasmid kit (Invitrogen). All sequences were confirmed by automated sequencing (Macrogen). The expression and purification of these antigens were the same as that of S-PCNA, M-PCNA2, H-PCNA3, or 6RBD-np.

### Size exclusion chromatography with multi-angle light scattering (SEC-MALS)

The shift in elution volume of the antigens was determined by size exclusion chromatography (SEC) on a Superdex 200 GL column (GE HealthCare) equilibrated with 50 mM Tris (pH 8.0) and 100 mM NaCl at a flow rate of 0.5 mL/min. To determine the molecular weight change of the antigens, purified protein (100 µL) was applied to Superdex 200 GL column (GE HealthCare) at 0.5 mL/min using a UFLC system (Shimadzu, Kyoto, Japan). Light scattering and refractive index were measured using in-line WYATT-787-TS miniDAWN TREOS (Wyatt Technology, Santa Barbara, CA, USA), which were analysed by Astra 6 software.

### Negative stained electron microscopy

Samples were prepared for electron microscopic studies by applying 6RBD-np (4 μL drop at a concentration of 1.5 μM) to charged EM grids coated with carbon film. Grids were negatively stained with 1% uranyl acetate and observed in a viewed under a Tecnai 20 transmission electron microscope operating at 120 kV (Thermo Fisher Scientific).

### Atomic force microscopy

The 6RBD-np (40 μL drop at 10 ng/mL) was deposited on a freshly cleaved mica surface and incubated for 15 min. To remove weakly bound protein residues and salts, the sample was washed with distilled water ten times and dried by N_2_ gas. The prepared sample was mounted on AFM scanner. The AFM data of 6RBD-np were acquired by soft tapping mode under ambient conditions using Multimode-Ⅷ (Bruker, Santa Barbara, CA, USA) with silicon nitride triangular AFM probes (SCANASYST-AIR, Bruker). The AFM images were captured at 512 × 512 pixels at 0.3 Hz in a line, and 2 nm in setpoint. The MountainsSPIP software (v9, Digital Surf, France) and Nansocope analysis software (v2.0, Bruker) were used to analyze the entire topographic information of 6RBD-np.

### Immunoblotting

For antigen characterization, purified antigens (RBD-SD1 of SARS-CoV-2, RBD-SD1 of SARS-CoV, S-PCNA1, M-PCNA2, H-PCNA3, and 6RBD-np) were subjected to immunoblotting, by incubating overnight with anti-6x His-tag polyclonal Ab (pAb) (ab1187) (AbCam plc, Cambridge, UK), rabbit anti-SARS-CoV-2 spike pAb (Sino Biological, 40592-T62), or anti-hCoV HKU1 spike monoclonal Ab (mAb) (40021-MM07-100, Sino Biological) diluted 1:1000 in TBS with 0.1% Tween 20.

### Mouse immunizations

Female BALB/c mice aged five weeks old were purchased from Koatech (Pyeongtaek, Kyunggi-do, Korea). Animal procedures were performed under the approval of the Institutional Animal Care and Use Committee (DSUA-2021-006-008) of Duksung Women’s University (Seoul, Korea). Experiments were done using mice at six weeks of age, with five mice for naive or PCNA control and ten mice for immunized groups: RBD-SD1 of SARS-CoV-2, S-2P trimer of SARS-CoV-2, S-PCNA1, and 6RBD-np. Mice were immunized in a prime-boost protocol with a three-week interval. Antigens were mixed with SAS adjuvant system (Sigma-Aldrich, St. Louis, MO, USA) to reach a final concentration of 0.05 mg/mL. Mice were injected intramuscularly into each hind leg with 50 μL per injection site (100 μL total). To obtain sera, mice were bled two weeks after prime and boost immunizations. Blood was collected via facial vein puncture in 1.5 mL Eppendorf tubes, which was stored at room temperature for 30 min to allow for coagulation. Serum was separated by centrifugation at 2,000 g for 15 min. Complement factors and pathogens in the serum were heat-inactivated at 56°C for 60 min, which were centrifuged and stored at −80°C until use.

### Antigen-specific Ab measurement by ELISA

Immunoglobulin G (IgG) responses were evaluated after prime and boost immunizations by enzyme-linked immunosorbent assay (ELISA). 96-well ELISA plates were coated with 100 μL of 2 μg/mL RBDs (SARS-CoV-2 WT and VAR, SARS-CoV, or hCoV HKU1) in PBS and incubated overnight at 4°C. The plates were washed with PBS after blocking with 1% bovine serum albumin (BSA) in PBS with 0.1% Tween 20 (PBS-T) for 1 hr at room temperature. 100 μL of serially diluted mouse sera, including naïve mouse serum, and blank (PBS-T) were added to the plate and incubated for 1 hr at room temperature. The plates were washed three times with PBS and dispensed with 100 μL of 1:5000 diluted goat anti-mouse IgG horseradish peroxidase for 1 hr at room temperature, followed by washing four times with PBS. 100 μL of 3,3′,5,5′-tetramethylbenzidine (Sigma-Aldrich) substrate were added to each well and incubated for 15 min at room temperature, which was stopped by adding 1 M H_2_SO_4_. The absorbance at a wavelength of 450 nm was measured using Multiskan™ FC Microplate Photometer (Thermo Fisher Scientific, Waltham, MA USA).

### Pseudovirus-based neutralization assays

Replication-deficient murine retrovirus-based pseudoviruses were generated by following a procedure previously described^52^. Briefly, site-directed mutagenesis and construction of four luciferase-expressing Maloney MLV-based pseudoviruses were carried out, encoding spike protein which represented SARS-CoV-2 Wuhan lineage (D614G), Delta lineage (L452R/T478K/D614G), two SD1 mutants (either N532A or E554A/N556A), or SARS-CoV, respectively. HEK293T cells were transfected with different plasmids encoding murine leukemia virus (MLV) gag/pol, SARS-CoV-2 spike D614G or other mutants, and firefly luciferase reporter. After 48 hrs post-transfection, pseudoviruses containing culture supernatants were harvested and filtered with 0.45-μm pore size (Millipore, Burlington, MA, USA). To quantify the titer of generated pseudoviruses, we carried out the quantitaive RT-PCR. Total RNA from the supernatant was extracted using Trisol solution (Thermo Fisher Scientific) and cDNA was synthesized with cDNA synthesis kit (Toyobo, Osaka, Japan). Real-time PCR was then performed with SYBR-green and CFX96 Real-Time System (Bio-rad, Carlsbad, CA, USA).

Pseudovirus containing spike protein was incubated with serial dilution of mouse sera for 30 min at room temperature, which was transduced into HEK293T-ACE2. After 24 hr transduction, cells were added with Bright-Glo (Promega), and relative luminescence units were measured by Varioskan Lux microplate reader (Thermo Fisher Scientific). All experiments were conducted in triplicates. The IC_50_ values were calculated with non-linear regression using GraphPad Prism 8 (GraphPad Software, Inc., San Diego, CA, USA).

### Live virus production

β-CoV/Korea/KCDC03/2020 (wild type: WT, S-clade), and hCoV-19/South Korea/KDCA2950/2021 (Delta variant) were propagated in Vero E6 cells grown in DMEM high glucose media (Gibco) supplemented with 10% FBS and 1% antibiotic-antimycotic in 37°C, 5% CO_2_.

### Virus challenge in immunized mice

To evaluate the immunogenicity and protective efficacy of the antigens in mice against the SARS-CoV-2 WT and Delta challenges, 6-week old female B6.Cg-Tg(K18-ACE2)2Prlmn/J mice were acclimatized for 7 days and given with intramuscular prime-boost immunization with a 3-week interval, 10 per dosing group per virus (WT and Delta) with 50 µL per injection site of vaccine formulations containing 5 µg of SARS-CoV-2 antigen (RBD-SD1, S-2P, S-PCNA1, or 6RBD-np) gently mixed 1:1 (v/v) with AddaVax (Invitrogen) adjuvant. The naïve control group received 50 µL PBS mixed 1:1 (v/v) with Addavax given intramuscularly. Animal experiments were approved by the Institutional Animal Care and Use Committee (CBNUA-1545-21-01) of Chungbuk National University (Chungbuk, Korea). Following the WHO recommendations, all work involving infectious SARS-CoV-2 was performed under biosafety level (BSL)-3 conditions in a facility with conditions at negative pressure and of operators in Tyvek suits wearing personal powered-air purifying respirators.

After 3 weeks from the second immunization, mice were inoculated with 10 MLD_50_ of SARS-CoV-2 WT or Delta variant intranasally under general anesthesia with isoflurane. Weight loss and survival rate were monitored for 14 days. All naïve animals did not lose weight and remained free of any disease signs throughout the study. Five days post-inoculation (dpi), lung and brain tissues were harvested from 3 mice per group and stored at −80°C until use for virus titration.

The collected frozen tissues were homogenized and serially diluted in DMEM. After homogenization of infected tissues, supernatant from tissue samples was 10-fold serially diluted and used to infect seeded monolayers of Vero E6 cells (1.5 x 10^4^/well) in 96-well plates. Infected cells were supplemented with DMEM containing 2% FBS and 1% antibiotic-antimycotic and incubated at 37°C, 5% CO_2_ for 96 hrs. The SARS-CoV-2 induced cytopathic effects (CPE) were measured using crystal violet staining. Viral titers in infected tissues were calculated using GraphPad Prism 9. Harvested lung samples at 5 dpi were fixed in 10% formalin neutralization buffer (Sigma-Aldrich, Saint Louis, MO, USA) and further processed for histopathological examination using hematoxylin-and-eosin (H&E) and immunohistochemical (IHC) staining.

### Pathology and immunohistochemistry

For histopathological examination, mouse lung samples harvested at 5 dpi were fixed in 10% formalin neutralization buffer (Sigma Aldrich) and stained with H&E. Additionally, IHC staining was performed to detect the viral antigen with primary antibodies, rabbit SARS-CoV-2 nucleocapsid pAb (40589-T62, Sinobiological, Beijing, China), at a dilution of 1:10,000 and HRP-conjugated secondary antibodies, using the Ventana discovery ULTRA system (Roche, Indianapolis, IN, USA).

### Virus-based neutralization assays

Sera from the blood of vaccinated mice were collected via the retro-orbital plexus at 14 and 35 days after prime immunization and stored at −80 °C until use. Vero E6 cells were prepared in a 96-well plate (1.5 x 10^4^ cells/well), and serum samples were diluted in two-fold dilution from 1:4 to 1:1024. SARS-CoV-2 viruses (WT or Delta variant) at 10^2^ TCID_50_/50 µl were prepared and mixed with diluted serum at a 1:1 ratio prior to 1 hr incubation at 37°C. Infected cells were supplemented with DMEM containing 2% FBS and 1% antibiotic-antimycotic and were incubated at 37°C, 5% CO_2_. After 96 hrs, SARS-CoV-2 induced CPE was measured using crystal violet staining and the serum neutralization titer (SN titer) was calculated

## Acknowledgments

This work was supported by the National Research Foundation of Korea grant (NRF-2021R1A4A1028969, K.H.K.) and the Korea Health Technology R&D Project grants (HV20C0054, K.H.K.) through the Korea Health Industry Development Institute (KHIDI), funded by the Korea government (MSIT and MHW) and the KBRI basic research program (21-BR-01-11, J.Y.M). The viruses, β-CoV/Korea/KCDC03/2020 (NCCP43326) and hCoV-19/South Korea/KDCA2950/2021 (NCCP43389), were provided by the National Culture Collection for Pathogens, Republic of Korea. We thank Dr. Giri Kotiguda for his initiation of a multivalent platform during an influenza vaccine development.

## Author Contributions

G.L., M.S., H.L., M.S.C., Y.H.B. and K.H.K. designed the study; D.B.L., H.K., J.H.J., E.S.J., E.J.K., S.Y.H., J.Y.M., S.M., M.R., M.S., H.L., M.S.C., Y.H.B., and K.H.K. prepared antigens, performed animal experiments and assays; S.R., H.J., J.Y.M, and G.L. performed TEM and AFM experiments; D.B.L., M.S., G.L., H.L., M.S.C., Y.H.B., and K.H.K. wrote the paper.

## Extended Data materials

**Extended Data Table 1.**
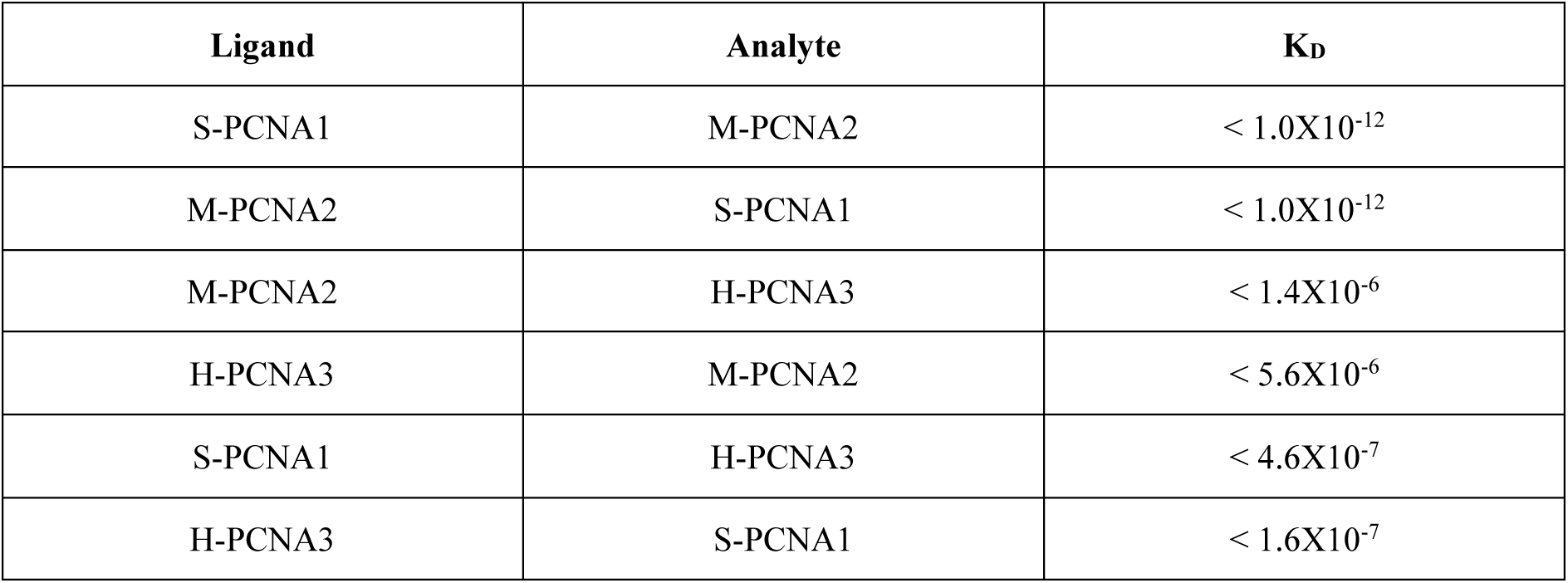
Dissociation constants (K_D_) of fusion proteins to assemble 6RBD-np.

**Extended Data Table 2.**
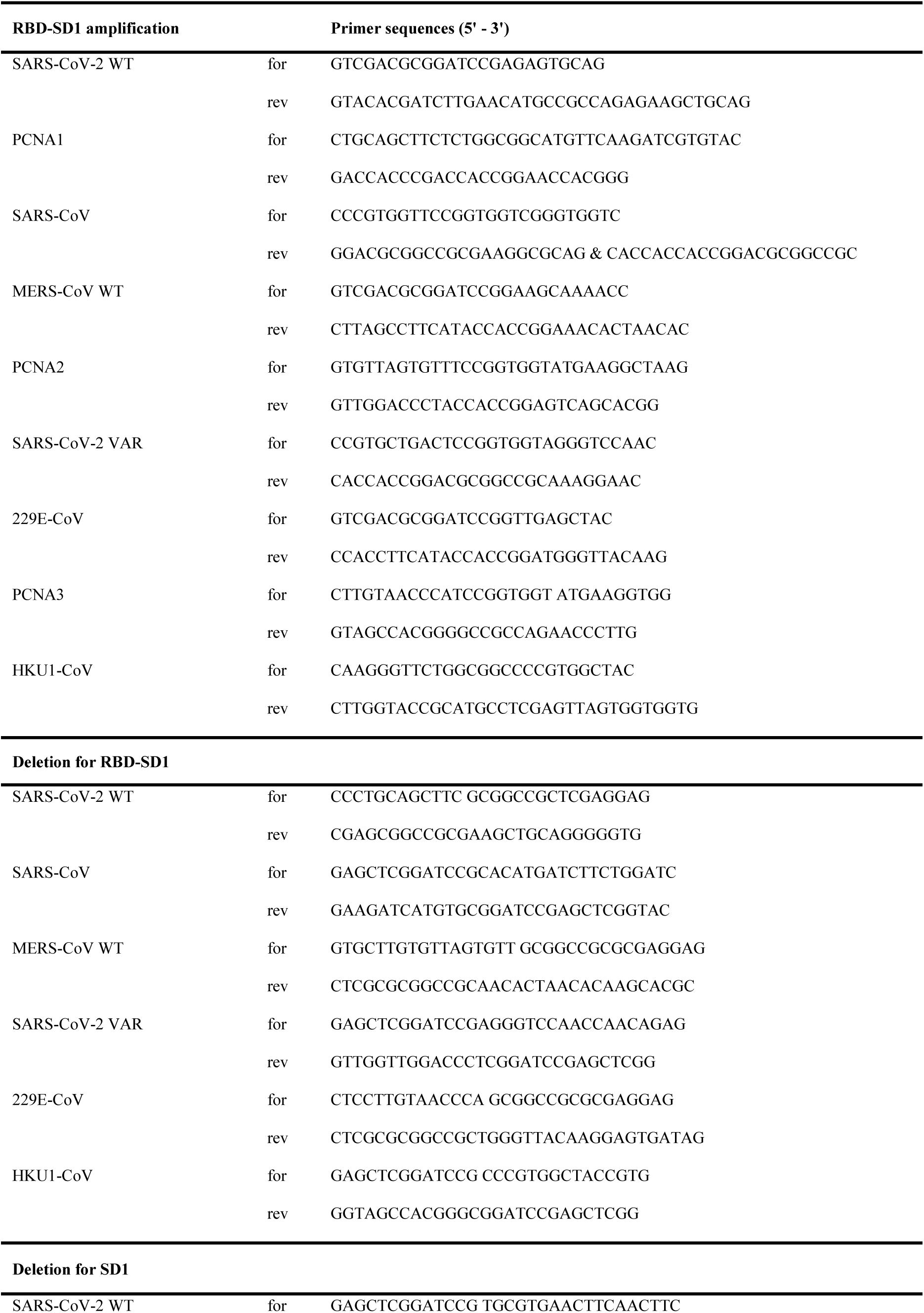
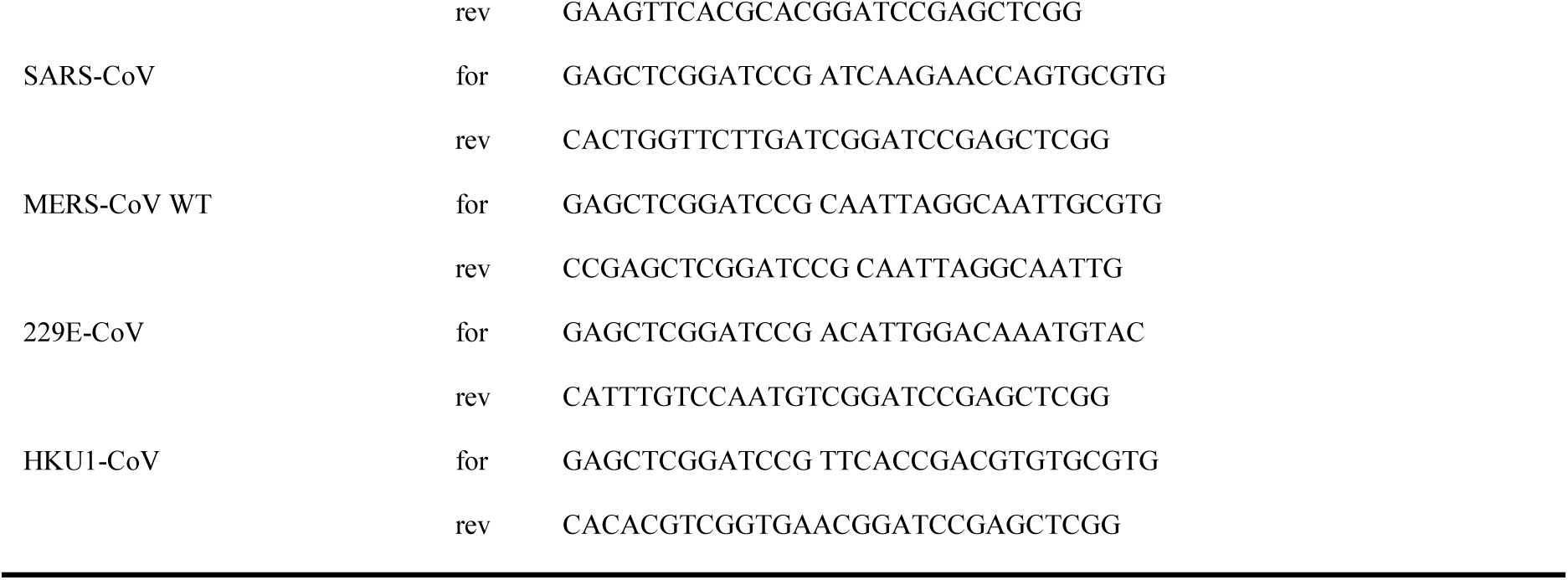
List of primers used in this study.

## Extended Data figure legends

**Extended Data Fig. 1.**
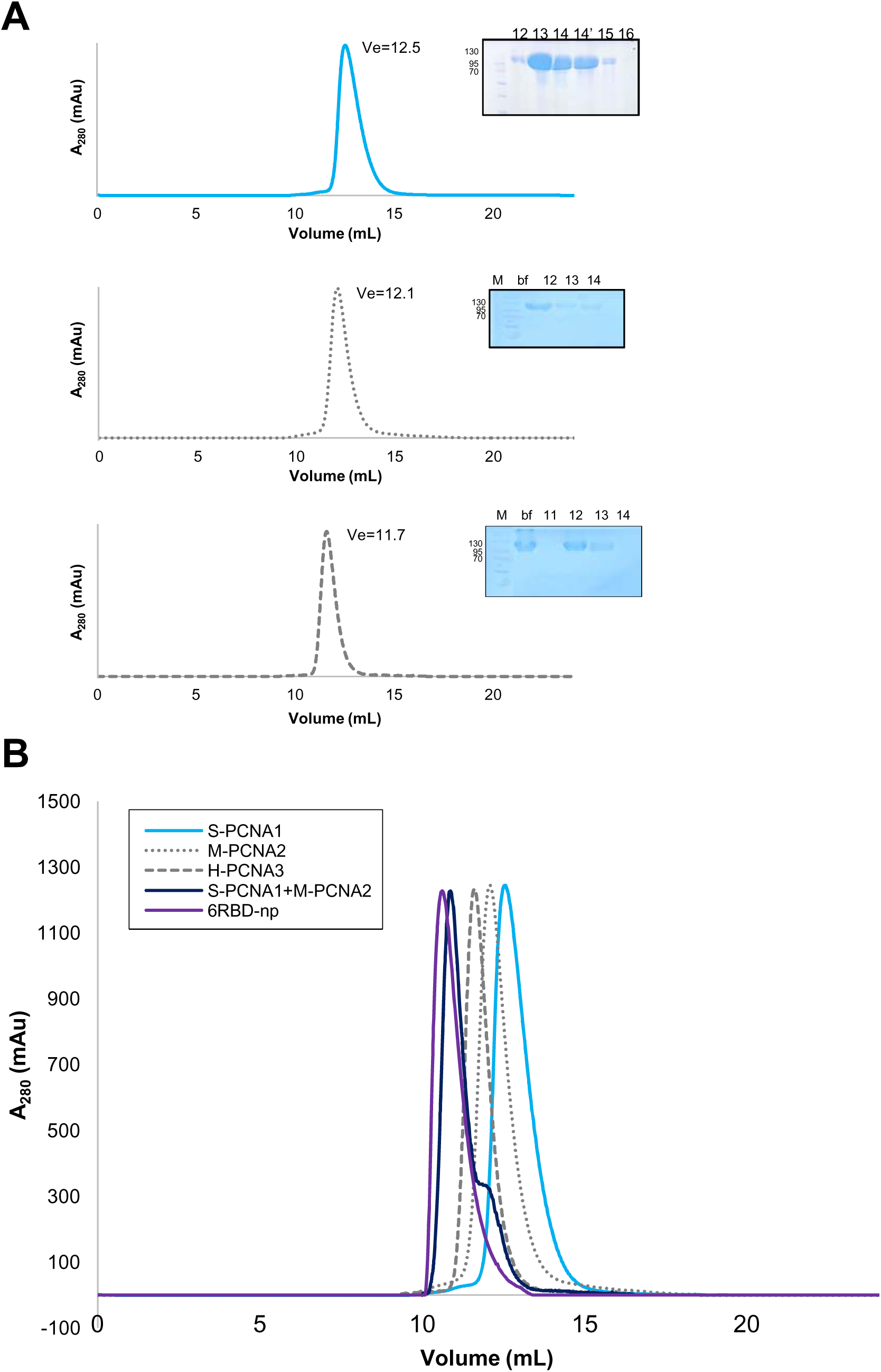
Purification of S-PCNA1, M-PCNA2 and H-PCNA3 and assembly of mosaic 6RBD-np. (A) Purification of S-PCNA1, M-PCNA2 and H-PCNA3. Their elution profiles from the final size exclusion chromatography steps, using Superdex 200 increase 10/300 GL, and SDS-PAGE results are shown on the right. The purified S-PCNA1, M-PCNA2 and H-PCNA3 have molecular weights in the range of 100−120 kDa, approximately consistent with their expected molecular weights of 97, 101, 120 kDa, respectively. (B) The shifts in elution profiles after assembly of 6RBD-np are shown for comparison, using size exclusion chromatography on Superdex 200 increase 10/300 GL. The purified S-PCNA1 and M-PCNA2 were mixed in a 1.2:1 molar ratio, to which H-PCNA3 was added in a molar ratio of 1.2:1.

**Extended Data Fig. 2.**
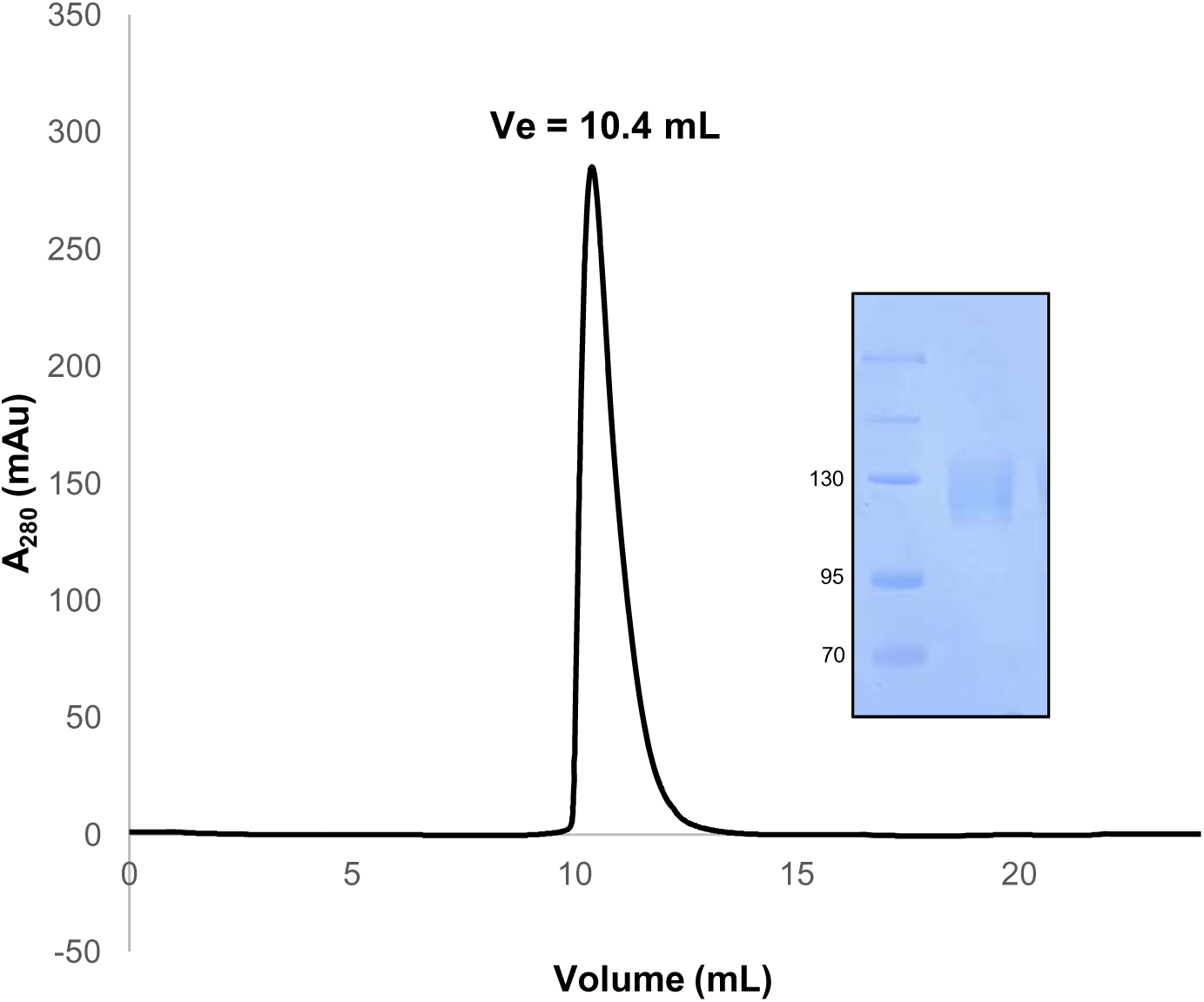
Characterization of 6RBD-np. The elution profile of 6RBD-np from the final size exclusion chromatography step, using Superdex 200 increase 10/300 GL, and SDS-PAGE results are shown on the right.

**Extended Data Fig. 3.**
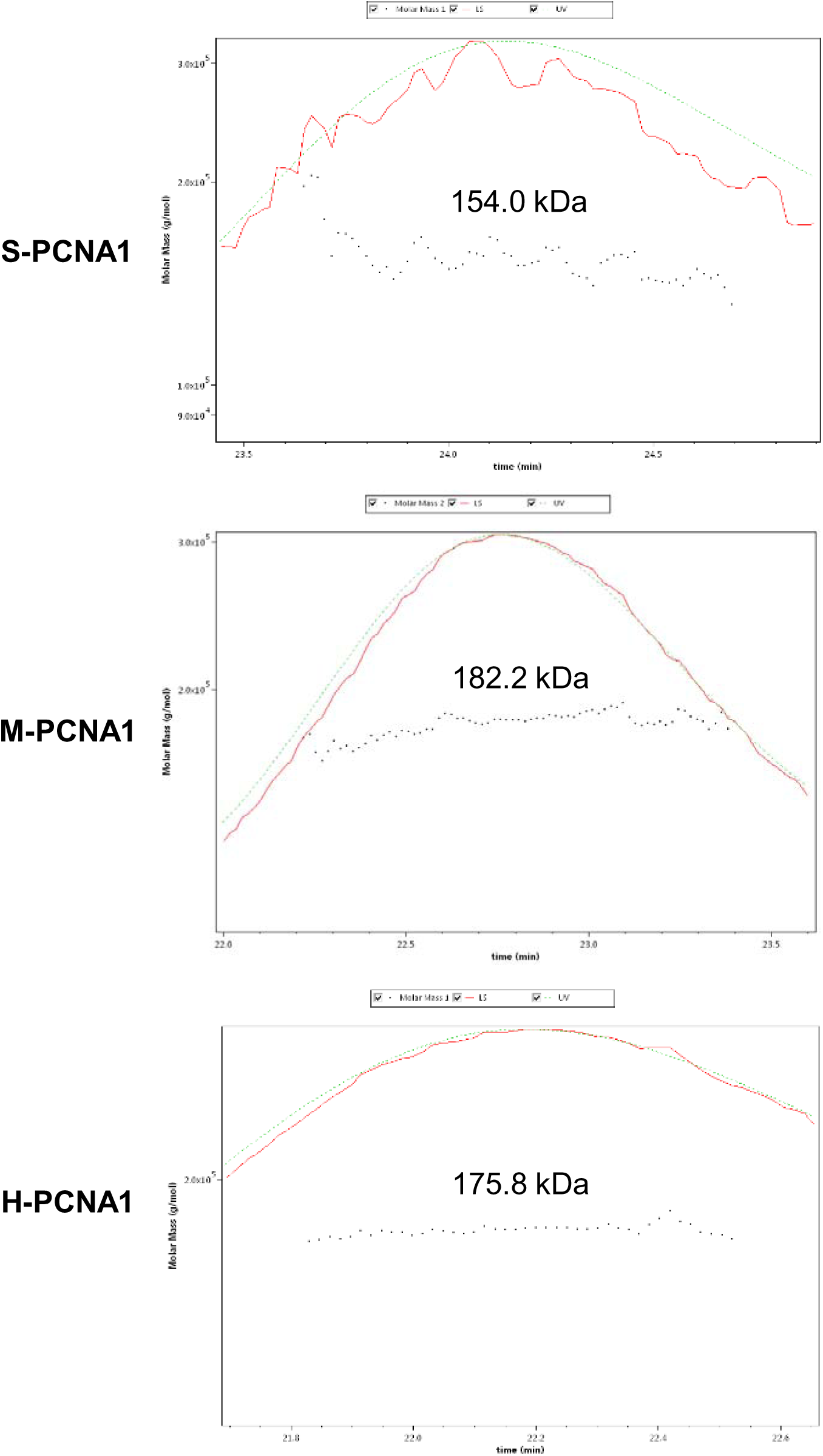
Characterization of 6RBD-np. The purified S-PCNA1, M-PCNA2 and H-PCNA3 are characterized using SEC-MALS to show molecular weights of 154.0 kDa (±12.2%), 182.2 kDa (±1.8%), and 175.8 kDa (±1.6%), respectively.

**Extended Data Fig. 4.**
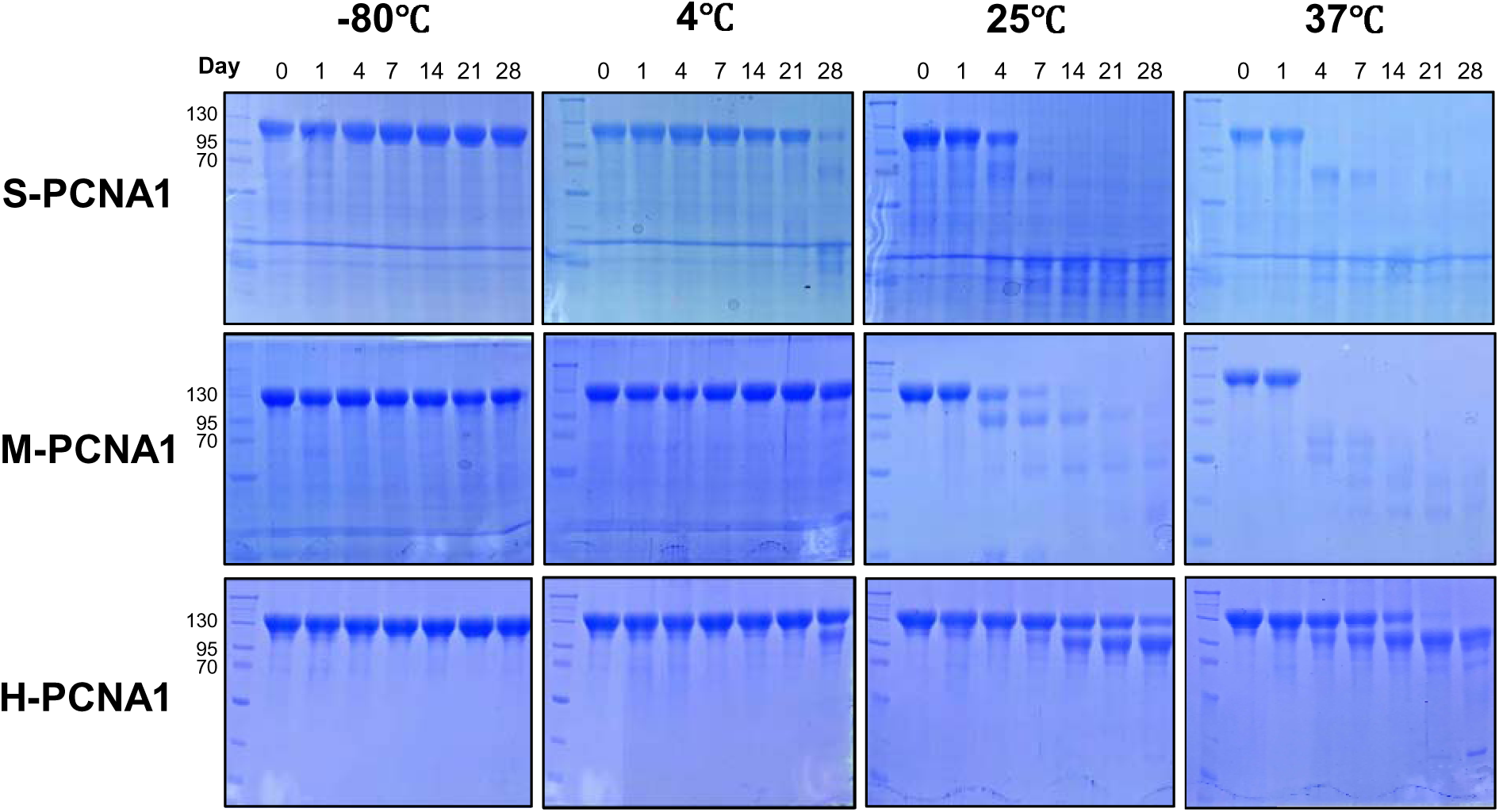
Stability of the 6RBD-np. Time-course experiments of S-PCNA1, M-PCNA2, and H-PCNA3 monitored at −80°C, 4°C, 25°C, and 37°C over a 28-day storage period using SDS-PAGE. The left lane, molecular weight marker, and lanes monitored on days 0, 1, 4, 7, 14, and 28.

**Extended Data Fig. 5.**
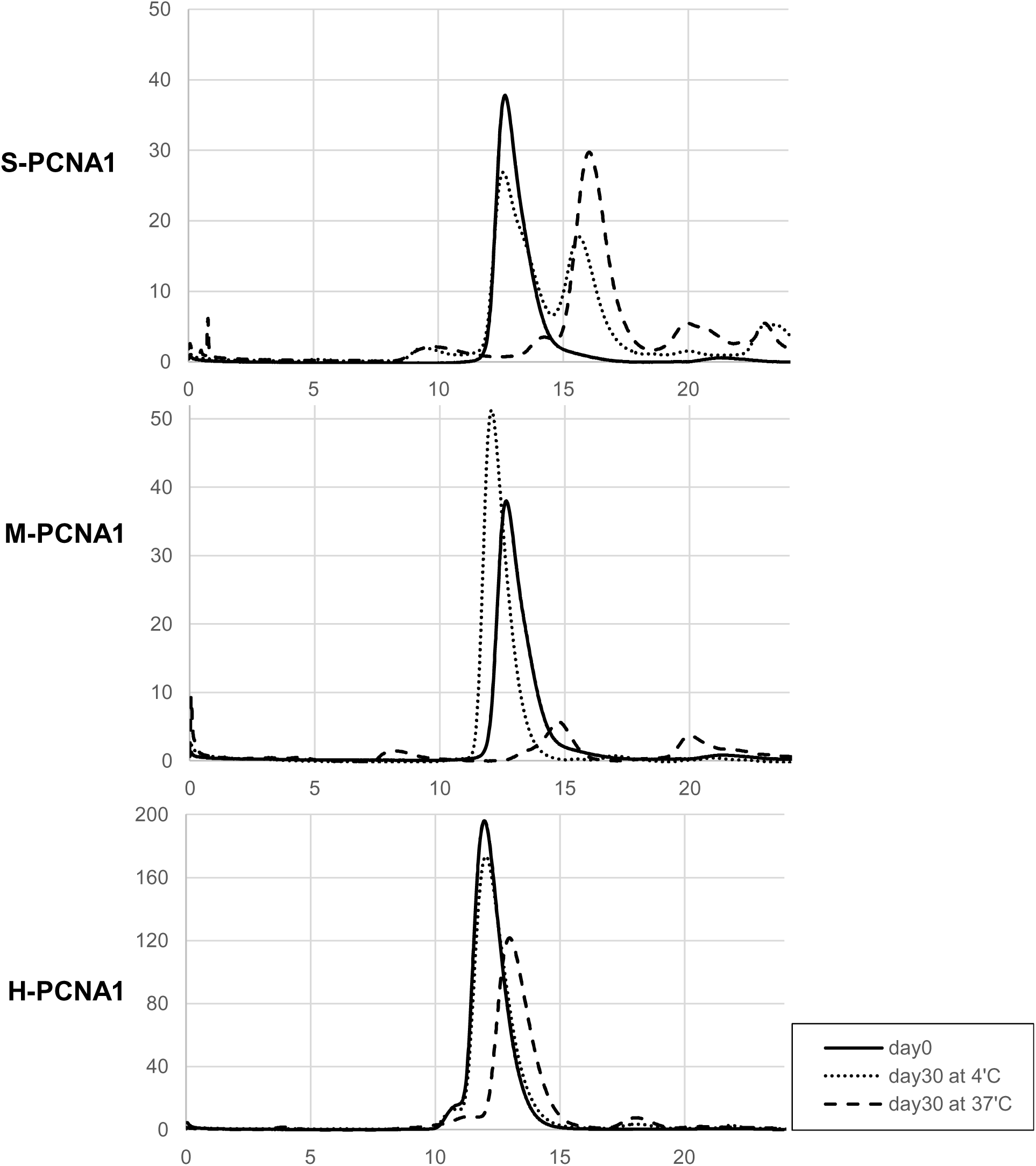
Stability of the 6RBD-np. Elution profiles of S-PCNA1, M-PCNA2 and H-PCNA3, obtained at day 0 (continuous line) and 30 at 4°C (dotted) and 37°C (dashed), after size exclusion chromatography on Superdex 200 increase 10/300 GL.

**Extended Data Fig. 6.**
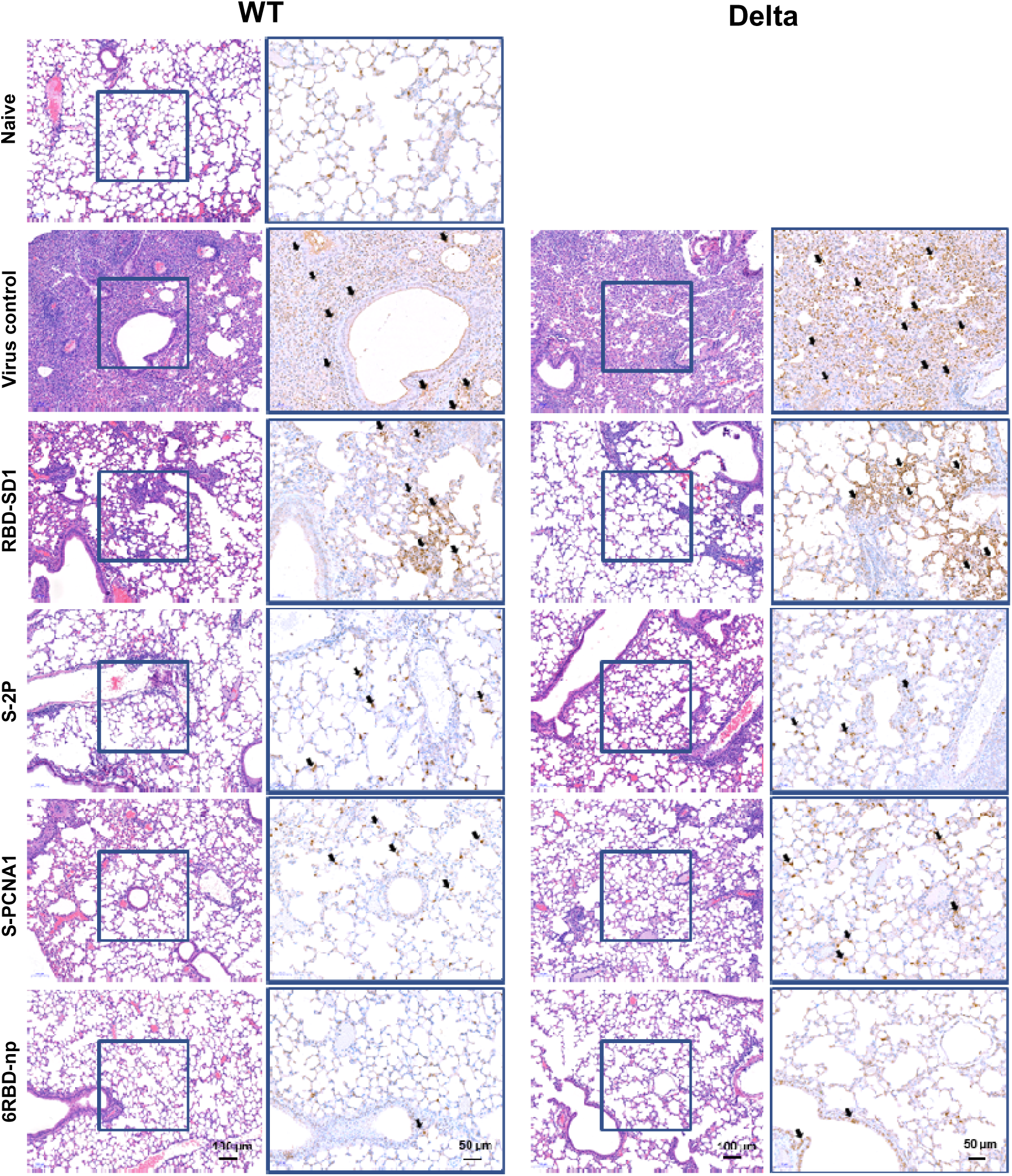
Representative images of H&E and IHC staining of lungs in mice infected with the wild type or Delta variant. Hematoxylin and eosin (scale bar, 100 μm) and immunohistochemistry (zoomed-in scale bar, 50 μm) staining of lung sections collected at 5 dpi for different groups are shown against SARS-CoV-2 wild type (left panel) and Delta variant (right panel) challenges. SARS-CoV-2 nucleoprotein (N) was detected in lung sections for IHC analysis in zoomed-in images of the box shown in the hematoxylin-and-eosin images, and the remainder was counterstained with hematoxylin. Arrows depicted in IHC staining indicate cells strongly detected by SARS-CoV-2 N protein specific antibody.

## References

1. Jones, K. E. et al. Global trends in emerging infectious diseases. Nature 451, 990–993, doi:10.1038/nature06536 (2008).

2. Taylor, L. H., Latham, S. M. & Woolhouse, M. E. Risk factors for human disease emergence. Philos Trans R Soc Lond B Biol Sci 356, 983–989, doi:10.1098/rstb.2001.0888 (2001).

3. Marston, H. D., Folkers, G. K., Morens, D. M. & Fauci, A. S. Emerging viral diseases: confronting threats with new technologies. Sci Transl Med 6, 253ps210, doi:10.1126/scitranslmed.3009872 (2014).

4. Cui, J., Li, F. & Shi, Z. L. Origin and evolution of pathogenic coronaviruses. Nat Rev Microbiol 17, 181–192, doi:10.1038/s41579-018-0118-9 (2019).

5. https://covid19.who.int.

6. Konings, F. et al. SARS-CoV-2 Variants of Interest and Concern naming scheme conducive for global discourse. Nat Microbiol 6, 821–823, doi:10.1038/s41564-021-00932-w (2021).

7. Viana, R. et al. Rapid epidemic expansion of the SARS-CoV-2 Omicron variant in southern Africa. 1–10 (2022).

8. Wang, Q. et al. Structural and functional basis of SARS-CoV-2 entry by using human ACE2. 181, 894–904. e899 (2020).

9. Greaney, A. J. et al. Antibodies elicited by mRNA-1273 vaccination bind more broadly to the receptor binding domain than do those from SARS-CoV-2 infection. Sci Transl Med 13, doi:10.1126/scitranslmed.abi9915 (2021).

10. Mishra, P. K. et al. Vol. 206 19 (The Journal of Immnology, 2021).

11. Jeyanathan, M. et al. Immunological considerations for COVID-19 vaccine strategies. 20, 615–632 (2020).

12. Krammer, F. A correlate of protection for SARS-CoV-2 vaccines is urgently needed. Nat Med 27, 1147–1148, doi:10.1038/s41591-021-01432-4 (2021).

13. Piccoli, L. et al. Mapping Neutralizing and Immunodominant Sites on the SARS-CoV-2 Spike Receptor-Binding Domain by Structure-Guided High-Resolution Serology. Cell 183, 1024–1042 e1021, doi:10.1016/j.cell.2020.09.037 (2020).

14. Yang, J. et al. A vaccine targeting the RBD of the S protein of SARS-CoV-2 induces protective immunity. Nature 586, 572–577, doi:10.1038/s41586-020-2599-8 (2020).

15. Dai, L. et al. A Universal Design of Betacoronavirus Vaccines against COVID-19, MERS, and SARS. Cell 182, 722–733 e711, doi:10.1016/j.cell.2020.06.035 (2020).

16. Yang, L. et al. A recombinant receptor-binding domain in trimeric form generates protective immunity against SARS-CoV-2 infection in nonhuman primates. Innovation (N Y) 2, 100140, doi:10.1016/j.xinn.2021.100140 (2021).

17. Walls, A. C. et al. Elicitation of Potent Neutralizing Antibody Responses by Designed Protein Nanoparticle Vaccines for SARS-CoV-2. Cell 183, 1367-+, doi:10.1016/j.cell.2020.10.043 (2020).

18. Kang, Y. F. et al. Rapid Development of SARS-CoV-2 Spike Protein Receptor-Binding Domain Self-Assembled Nanoparticle Vaccine Candidates. ACS Nano 15, 2738–2752, doi:10.1021/acsnano.0c08379 (2021).

19. Saunders, K. O. et al. Neutralizing antibody vaccine for pandemic and pre-emergent coronaviruses. Nature 594, 553–559, doi:10.1038/s41586-021-03594-0 (2021).

20. Wall, E. C. et al. Neutralising antibody activity against SARS-CoV-2 VOCs B. 1.617. 2 and B. 1.351 by BNT162b2 vaccination. 397, 2331–2333 (2021).

21. Goldberg, Y. et al. Waning Immunity after the BNT162b2 Vaccine in Israel. N Engl J Med 385, e85, doi:10.1056/NEJMoa2114228 (2021).

22. Mizrahi, B. et al. Correlation of SARS-CoV-2-breakthrough infections to time-from-vaccine. Nat Commun 12, 6379, doi:10.1038/s41467-021-26672-3 (2021).

23. Collier, D. A. et al. Sensitivity of SARS-CoV-2 B.1.1.7 to mRNA vaccine-elicited antibodies. Nature 593, 136–141, doi:10.1038/s41586-021-03412-7 (2021).

24. McCallum, M. et al. SARS-CoV-2 immune evasion by the B.1.427/B.1.429 variant of concern. Science 373, 648–654, doi:10.1126/science.abi7994 (2021).

25. Wang, P. F. et al. Antibody resistance of SARS-CoV-2 variants B.1.351 and B.1.1.7. Nature 593, 130–135, doi:10.1038/s41586-021-03398-2 (2021).

26. Kanekiyo, M. et al. Mosaic nanoparticle display of diverse influenza virus hemagglutinins elicits broad B cell responses. Nature Immunology 20, 362–372 (2019).

27. Walls, A. C. et al. Elicitation of broadly protective sarbecovirus immunity by receptor-binding domain nanoparticle vaccines. Cell 184, 5432–5447, doi:10.1016/j.cell.2021.09.015 (2021).

28. Cohen, A. A. et al. Mosaic nanoparticles elicit cross-reactive immune responses to zoonotic coronaviruses in mice. Science 371, 735–741, doi:10.1126/science.abf6840 (2021).

29. Dionne, I., Nookala, R. K., Jackson, S. P., Doherty, A. J. & Bell, S. D. A heterotrimeric PCNA in the hyperthermophilic archaeon Sulfolobus solfataricus. Mol Cell 11, 275–282, doi:10.1016/s1097-2765(02)00824-9 (2003).

30. Williams, G. J. et al. Structure of the heterotrimeric PCNA from Sulfolobus solfataricus. Acta Crystallogr Sect F Struct Biol Cryst Commun 62, 944–948, doi:10.1107/S1744309106034075 (2006).

31. Hlinkova, V. et al. Structures of monomeric, dimeric and trimeric PCNA: PCNA-ring assembly and opening. Acta Crystallogr D Biol Crystallogr 64, 941–949, doi:10.1107/S0907444908021665 (2008).

32. Tan, C. K., Castillo, C., So, A. G. & Downey, K. M. An auxiliary protein for DNA polymerase-delta from fetal calf thymus. J Biol Chem 261, 12310–12316 (1986).

33. Liu, L. et al. Potent neutralizing antibodies against multiple epitopes on SARS-CoV-2 spike. Nature 584, 450–456, doi:10.1038/s41586-020-2571-7 (2020).

34. Brouwer, P. J. M. et al. Potent neutralizing antibodies from COVID-19 patients define multiple targets of vulnerability. Science 369, 643–650, doi:10.1126/science.abc5902 (2020).

35. Robbiani, D. F. et al. Convergent antibody responses to SARS-CoV-2 in convalescent individuals. Nature 584, 437–442, doi:10.1038/s41586-020-2456-9 (2020).

36. Sanders, R. W. & Moore, J. P. J. I. r. Native-like Env trimers as a platform for HIV-1 vaccine design. Immunol Rev. 275, 161–182 (2017).

37. Khoury, D. S. et al. Neutralizing antibody levels are highly predictive of immune protection from symptomatic SARS-CoV-2 infection. Nat Med 27, 1205–1211, doi:10.1038/s41591-021-01377-8 (2021).

38. Wang, R., Chen, J., Gao, K. & Wei, G. W. Vaccine-escape and fast-growing mutations in the United Kingdom, the United States, Singapore, Spain, India, and other COVID-19-devastated countries. Genomics 113, 2158–2170, doi:10.1016/j.ygeno.2021.05.006 (2021).

39. Tian, J. H. et al. SARS-CoV-2 spike glycoprotein vaccine candidate NVX-CoV2373 immunogenicity in baboons and protection in mice. Nat Commun 12, 372, doi:10.1038/s41467-020-20653-8 (2021).

40. Polack, F. P. et al. Safety and Efficacy of the BNT162b2 mRNA Covid-19 Vaccine. N Engl J Med 383, 2603–2615, doi:10.1056/NEJMoa2034577 (2020).

41. Baden, L. R. et al. Efficacy and Safety of the mRNA-1273 SARS-CoV-2 Vaccine. N Engl J Med 384, 403–416, doi:10.1056/NEJMoa2035389 (2021).

42. Hillus, D. et al. Safety, reactogenicity, and immunogenicity of homologous and heterologous prime-boost immunisation with ChAdOx1-nCoV19 and BNT162b2: a prospective cohort study. Lancet Respir Med. (2021).

43. Pulliam, J. R., et al. Increased risk of SARS-CoV-2 reinfection associated with emergence of the Omicron variant in South Africa. MedRxiv (2021).

44. Pajon, R. et al. SARS-CoV-2 Omicron Variant Neutralization after mRNA-1273 Booster Vaccination. N Engl J Med, doi:10.1056/NEJMc2119912 (2022).

45. McCallum, M. et al. Structural basis of SARS-CoV-2 Omicron immune evasion and receptor engagement. Science, eabn8652, doi:10.1126/science.abn8652 (2022).

46. Bachmann, M. F. et al. The influence of antigen organization on B cell responsiveness. Science 262, 1448–1451, doi:10.1126/science.8248784 (1993).

47. Wuertz, K. M. et al. A SARS-CoV-2 spike ferritin nanoparticle vaccine protects hamsters against Alpha and Beta virus variant challenge. NPJ Vaccines 6, 129, doi:10.1038/s41541-021-00392-7 (2021).

48. Nguyen, B. & Tolia, N. H. Protein-based antigen presentation platforms for nanoparticle vaccines. NPJ Vaccines 6, 70, doi:10.1038/s41541-021-00330-7 (2021).

49. Bachmann, M. F. & Jennings, G. T. Vaccine delivery: a matter of size, geometry, kinetics and molecular patterns. Nat Rev Immunol 10, 787–796, doi:10.1038/nri2868 (2010).

50. FDA, U. J. S. S., MD: US Department of Health & Services, H. Drug Products, Including Biological Products, That Contain Nanomaterials—Guidance for Industry. (2017).

51. Jegerlehner, A. et al. Regulation of IgG antibody responses by epitope density and CD21- mediated costimulation. Eur J Immunol 32, 3305–3314, doi:10.1002/1521-4141(200211)32:11<3305::AID-IMMU3305>3.0.CO;2-J (2002).

52. Kim, Y. J., Jang, U. S., Soh, S. M., Lee, J. Y. & Lee, H. R. The Impact on Infectivity and Neutralization Efficiency of SARS-CoV-2 Lineage B.1.351 Pseudovirus. Viruses 13, 633, doi:10.3390/v13040633 (2021).

